# ABA-regulated JAZ1 Proteins Bind NAC42 Transcription Factors to Suppress the Activation of Phytoalexin Biosynthesis in Plants

**DOI:** 10.1101/2024.09.26.615281

**Authors:** Jie Lin, Ivan Monsalvo, Md Asraful Jahan, Melissa Ly, Dasol Wi, Izabella Martirosyan, Israt Jahan, Nik Kovinich

## Abstract

Phytoalexins are plant defense metabolites whose biosynthesis remains suppressed until elicited by a pathogen or stress, yet the mechanism of their suppression has remained elusive. The transcription factor GmNAC42-1 is an important and direct activator of the biosynthesis of glyceollin phytoalexins in soybean. Yet, without elicitation, overexpressing GmNAC42-1 is insufficient to activate the expression of glyceollin biosynthetic genes, suggesting that the activity of GmNAC42-1 may be suppressed by a negative regulator. JAZ1 proteins are negative regulators of the canonical jasmonic acid (JA) signaling pathway. JAZ protein degradation and JAZ gene transcription comprise antagonistic mechanisms that activate and suppress JA signaling, respectively. In search for negative regulators of glyceollin biosynthesis, we identified by RNA-seq analysis abscisic acid (ABA) signaling and GmJAZ1 genes that are oppositely regulated compared to glyceollin biosynthesis. Long-term ABA treatment upregulated GmJAZ1 transcripts, whereas its biosynthesis inhibitor fully suppressed their upregulation by dehydration stress. Opposite patterns were observed for glyceollin biosynthesis. RNAi silencing of GmJAZ1s prevented the suppression of glyceollin biosynthesis by dehydration and derepressed glyceollin synthesis in non-elicited tissues. Overexpressing GmJAZ1-9 in hairy roots elicited with Phytophthora sojae wall glucan elicitor partially suppressed glyceollin biosynthesis. The GmJAZ1-9 protein physically interacted with GmNAC42-1 and inhibited its transactivation and DNA binding activities in promoter-luciferase and yeast-three hybrid systems. Silencing JAZ1s in Arabidopsis and grapevine has been reported to derepress camalexin and stilbene phytoalexin biosynthesis. Here, we found that JAZ1 and NAC42 proteins from all three plant species physically interact, suggesting a conserved mechanism negatively regulates phytoalexin biosynthesis in plants.

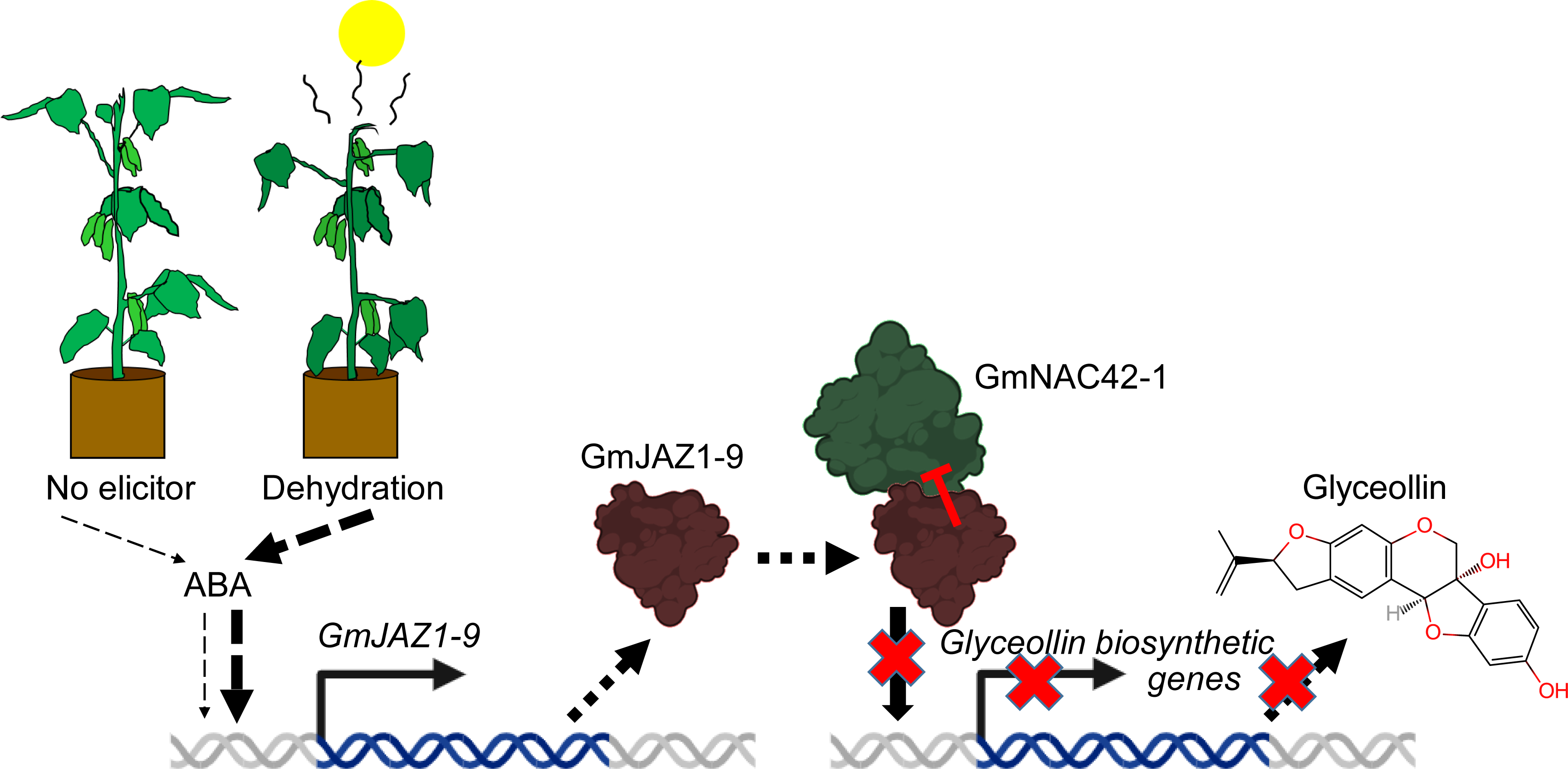

## INTRODUCTION

Phytoalexins are plant specialized metabolites that are biosynthesized in response to pathogens and particular abiotic stresses. Genetic and biochemical evidence suggests that phytoalexins play pivotal roles in mediating incompatibility between plants and pathogens (Ahuja *et al*., 2012, Großkinsky *et al*., 2012, Jeandet *et al*., 2014). Silencing *GmMYB29A2*, a transcription factor gene that positively regulates glyceollin biosynthesis in soybean, renders the R gene *Rps1k* variety Williams 82 compatible with Race 1 *Phytophthora sojae* (Jahan *et al*., 2020). Conversely, overexpressing *GmMYB29A2* in the susceptible variety Williams, that lacks *Rps1k*, renders it incompatible with Race 1 *P. sojae*. Ectopically overexpressing grapevine (*Vitis vinifera*) stilbene synthase (*STS*) genes in heterologous plant hosts provides resistance to several necrotrophic and hemibiotrophic pathogens (Chong *et al*., 2009, Zhang *et al*., 2023). Also, mutants of Arabidopsis (*Arabidopsis thaliana*) that are deficient in camalexin biosynthesis, or its regulation, are susceptible to a wide variety of necrotrophic, hemibiotrophic, and biotrophic pathogens (Nguyen *et al*., 2022). Phytoalexin biosynthesis and its regulation is achieved by complex gene networks, the study of which continues to inform on fundamental aspects of plant biology (Zhao *et al*., 2021, Polturak *et al*., 2022, Zhou *et al*., 2022). Understanding the transcription factor networks that regulate phytoalexin biosynthesis could lead to enhanced phytoalexin production via biotechnology or breeding efforts. It could improve the production of phytoalexins that are used as clinical pharmaceuticals, including taxol, berberine, and artemisinin (Dittrich and Kutchan, 1991, Li *et al*., 2012, Chen *et al*., 2021). It also could improve accessibility to phytoalexins that exhibit promising properties for pharmaceutical development (Egbuonu and Eneogwe, 2018, Ahmed and Kovinich, 2021, Sivakumar and Deepa, 2023). These include the neuroprotective activities of glyceollins, stilbenes, and camalexin (Seo *et al*., 2018, Rahman *et al*., 2020, Thamaraikani *et al*., 2022). Glyceollins exhibit unconventional anti-tumor activities against breast and lung cancer types that are resistant to conventional therapeutics (Rhodes *et al*., 2012, Lee *et al*., 2015, Walker *et al*., 2022). Pterostilbene inhibits the invasion, migration, and metastasis of human hepatoma cells (Obrador *et al*., 2021), and camalexin selectively inhibits the proliferation of prostate cancer over non-cancerous cells (Pilatova *et al*., 2013). Since plants remain the most economical source of many phytoalexins, understanding phytoalexin biosynthesis and its regulation remain important goals for agricultural and pharmaceutical sciences (Egbuonu and Eneogwe, 2018, Mattio *et al*., 2020, Sharma *et al*., 2022).

Phytoalexins are biosynthesized from diverse metabolic pathways that generally differ among plant lineages (Wu, 2020, Bizuneh, 2021, Largia *et al*., 2023). For example, glyceollins are biosynthesized from phenylalanine *via* the isoflavonoid pathway, stilbenes are from tyrosine or phenylalanine *via* the phenylpropanoid pathway, and camalexin is biosynthesized from tryptophan *via* the indole alkaloid pathway (Figure 1). Despite their biosynthetic heterogeneity, we have recently discovered that the same conserved transcription factors activate camalexin, stilbene, and glyceollin biosynthesis in Arabidopsis, grapevine, and soybean, respectively (Jahan *et al*., 2019, Jahan *et al*., 2020). The soybean NAC family transcription factor GmNAC42-1 directly binds the promoter DNA of the glyceollin biosynthetic genes *IFS2* and *G4DT* in the yeast one-hybrid (Y1H) system (Jahan *et al*., 2019). *GmNAC42-1* is the homolog of *ANAC042*, which is an important regulator of camalexin biosynthesis in Arabidopsis (Saga *et al*., 2012). *anac042* loss-of-function mutants have reduced expression of the camalexin biosynthetic genes *CYP71A12*, *CYP71A13*, and *CYP71B15/PAD3*, yet it remains to be determined whether ANAC042 directly regulates those genes. GmMYB29A2 is an R2R3 MYB family transcription factor that directly binds the promoters of the glyceollin biosynthetic genes *IFS2* and *G4DT* in the Y1H system and in electrophoretic mobility shift assays (EMSAs) (Jahan *et al*., 2020). *GmMYB29A2* is the soybean homolog of *VvMYB14/VvMYB15*, which encode transcription factors that directly activate stilbene synthase (*STS*) genes in grapevine (Höll *et al*., 2013, Orduña *et al*., 2022). The homolog in Arabidopsis, namely *MYB15*, activates biosynthesis of scopoletin and pathogen-inducible monolignol phytoalexins (Chezem *et al*., 2017). RNAi silencing demonstrated that both *GmMYB29A2* and *GmNAC42-1* are required for activating glyceollin biosynthesis in soybean in response to *P. sojae* WGE (Jahan *et al*., 2019, Jahan *et al*., 2020). Yet, co-overexpressing *GmMYB29A2* and *GmNAC42-1* in the absence of an elicitor treatment was insufficient to activate glyceollin biosynthesis (Lin et al., 2023a). Thus, we hypothesized that one or more negative regulators remain unidentified that are needed to fully unlock the regulation of phytoalexin biosynthesis. In our search for genes that are co-expressed with glyceollin biosynthesis, we recently identified the WRKY family transcription factor GmWRKY72 (Lin *et al*., 2023b, Lin *et al*., 2024). Overexpressing and silencing its gene in the soybean hairy root system found that GmWRKY72 is a negative regulator of glyceollin biosynthesis that is active only in elicited cells (Lin *et al*., 2024). In non-elicited tissues, RNAi silencing of GmWRKY72 did not result in any increases in glyceollin amounts. Thus, the mechanism that keeps glyceollin biosynthesis ‘off’ in the absence of an elicitor remained unknown. Our recent review of phytoalexin gene regulation in Arabidopsis and its comparison to crop plants suggested that the transcription factors ANAC042, MYB15, WRKY33, ERF1, and MYB72 may comprise a ‘core’ transcription factor network for the positive regulation of phytoalexin biosynthesis in plants, and JAZ1 proteins may represent a conserved negative regulator of that network (Monsalvo *et al*., 2024).

**Figure 1.**
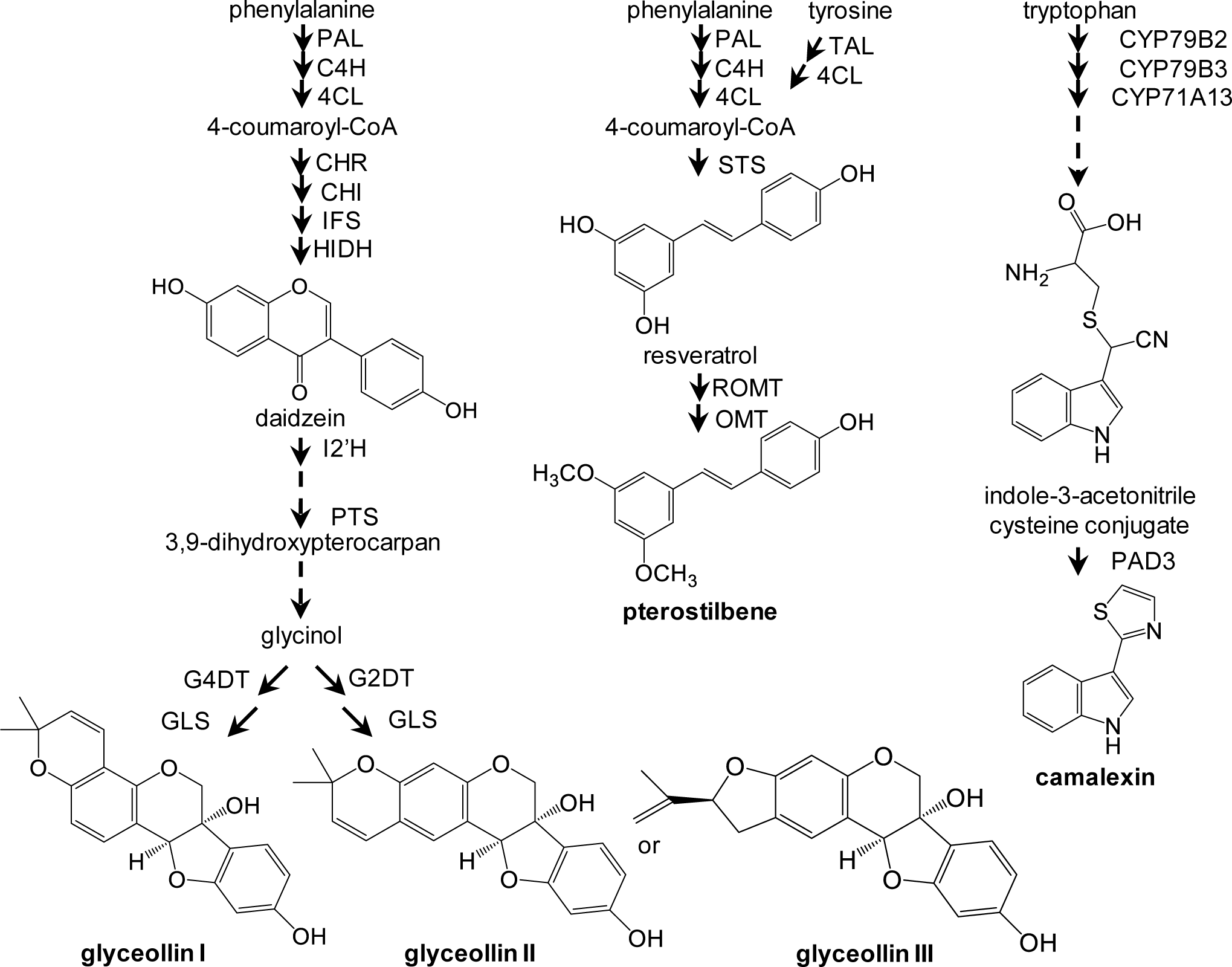
Soybean glyceollin, grapevine stilbene, and Arabidopsis camalexin biosynthetic pathways. Solid and broken arrows indicate single and multiple enzymatic steps, respectively. PAL, phenylalanine ammonia lyase; C4H, cinnamate-4-hydroxylase; 4CL, 4-Coumarate-CoA ligase; CHR, chalcone reductase; CHI, chalcone isomerase; IFS, isoflavone synthase; HIDH, 2−hydroxyisoflavanone dehydratase; I2’H, isoflavone 2′-hydroxylase; PTS, pterocarpan synthase; G4DT, glycinol 4-dimethylallyl transferase; G2DT, glycinol 2-dimethylallyl transferase; GLS, glyceollin synthase; TAL, tyrosine ammonia lyase; STS, stilbene synthase; ROMT, resveratrol *O*-methyltransferase; OMT, *O*-methyltransferase; CYP, cytochrome P450; PAD3, phytoalexin deficient 3 (*a.k.a.* CYP71B15).

Applying the plant hormone abscisic acid (ABA) to plant tissues negatively regulates phytoalexin biosynthesis in various plant species (Henfling *et al*., 1981, Ward *et al*., 1989). In soybean upon WGE treatment, endogenous ABA levels decrease prior to activating the transcription of glyceollin biosynthetic genes (Mohr *et al*., 2001). In tobacco, the enzymatic deactivation of ABA to 8’-hydroxy-ABA coincides with the activation of capsidiol phytoalexin biosynthesis (Mialoundama *et al*., 2009). In Arabidopsis, the pathogen *Pseudomonas syringae* exploits ABA-induced protein phosphatase 2Cs (PP2Cs), HAI1, HAI2, and HAI3, to dephosphorylate the protein kinases MPK3 and MPK6 (Mine *et al*., 2017). Thus, ABA signaling may be a conserved suppressor of phytoalexin biosynthesis, but its mechanism of negative regulation remains unknown. Several transcription factors and proteins have been identified that negatively regulate phytoalexin biosynthesis. VvWRKY8 physically interacts with VvMYB14 to prevent its binding to DNA and its transactivation of *STS* genes (Jiang *et al*., 2019). The paralog of *GmMYB29A2*, namely *GmMYB29A1*, reduces glyceollin accumulation when overexpressed in soybean hairy roots without affecting the expression of most glyceollin biosynthetic genes (Jahan *et al*., 2020). In Arabidopsis, the ubiquitin E3 ligase SR1 degrades phosphorylated WRKY33, which is an important transcriptional activator of camalexin biosynthesis (Liu *et al*., 2021). Further, RNAi silencing of *JAZ1* in Arabidopsis calli, or its homolog *VvJAZ9* in grapevine cell cultures, derepresses camalexin and stilbene phytoalexin biosynthesis, respectively (Makhazen *et al*., 2021a, Makhazen *et al*., 2021b). JAZ proteins are negative regulators of the JA signaling pathway (Chini *et al*., 2007). JAZ proteins are components of the COI1-JAZ co-receptor that binds and mediates the perception of the bioactive form of JA, namely jasmonoyl-isoleucine (JA-Ile) (Sheard *et al*., 2010). JAZ proteins negatively regulate the JA signaling pathway by physically binding transcription factor proteins to inhibit their activities (Cheng *et al*., 2011, Fernández-Calvo *et al*., 2011, Qi *et al*., 2011, Sasaki-Sekimoto *et al*., 2014). Upon elicitation, JA-Ile is biosynthesized and binds to the COI1-JAZ co-receptor to stimulate the 26S proteasome-mediated degradation of JAZ proteins (Chini *et al*., 2007). This removes JAZ proteins from their interacting transcription factors allowing those factors to activate the transcription of downstream genes. JAZ protein degradation activates JA signaling, yet JA signaling positively regulates the transcription of *JAZ* genes. This has been proposed to result in a pulsed response, whereby JA signaling is activated, then repressed *via* the upregulation of *JAZ* gene transcription (Chini *et al*., 2007). Thus, the balance between JAZ protein degradation and *JAZ* gene transcription are important for determining the level of activation or repression of JA signaling. Pathways that positively regulate *JAZ* gene transcription remain largely unknown.

The transcriptional regulation of phytoalexins is a topic of great interest because of their potential utility in developing disease-resistant crops and plant-based platforms for pharmaceutical production. Here, we found that the suppression of glyceollin biosynthesis in soybean by long-term dehydration stress is dependent ABA signaling and its transcriptionally upregulated *GmJAZ1* genes. JAZ1 proteins from several plant species physically interact with NAC42-type transcriptional activators of phytoalexin biosynthesis. The derepression of phytoalexin biosynthesis by silencing JAZ1 genes may be a vital step towards engineering or breeding plants for the stable supply of valuable phytoalexin molecules.

## RESULTS

### RNA-seq identifies ABA signaling and *GmJAZ1* genes as candidate negative regulators of glyceollin biosynthesis

We have previously found that dehydration stress suppresses and acidic (pH 3.0) medium enhances the elicitation of glyceollin biosynthesis (Jahan *et al*., 2019). These stresses reflect drought conditions, and extreme soil pH conditions such as that of mine tailing sites, respectively. To search for candidate negative regulators of glyceollin biosynthesis, we analyzed our RNA-seq data of seedlings grown in those two stress conditions and identified 731 genes that had opposite expression patterns compared to glyceollin biosynthetic genes (Figure 2A). The majority of those genes (16.1%) were categorized as signaling genes, of which 20.5% were annotated as ABA genes and 19.2% as JA genes (Figure 2B, 2C). Liquid chromatography-mass spectrometry (LC-MS) analysis found that dehydration-stressed seedlings accumulated the metabolite ABA but had reduced amounts of JA (Figure 2D). In contrast, pH 3.0 medium induced the accumulation of the storage conjugate of ABA, namely ABA-glucose ester (ABA-GE), and had elevated levels of JA. A repeat of this experiment demonstrated similar results (Supplementary Figure S1). Statistical values for these experiments are shown in Supplementary Table S1. The expression of six JASMONATE-ZIM-DOMAIN 1 (GmJAZ1) homologs and several ABA marker genes, such as *ATP-BINDING CASSETTE TRANSPORTER G40* and *RESPONSIVE TO DESICCATION 22* (*GmRD22*), were upregulated by dehydration stress, similar to ABA metabolite profiles (Table 1, Figure 2D).

**Figure 2.**
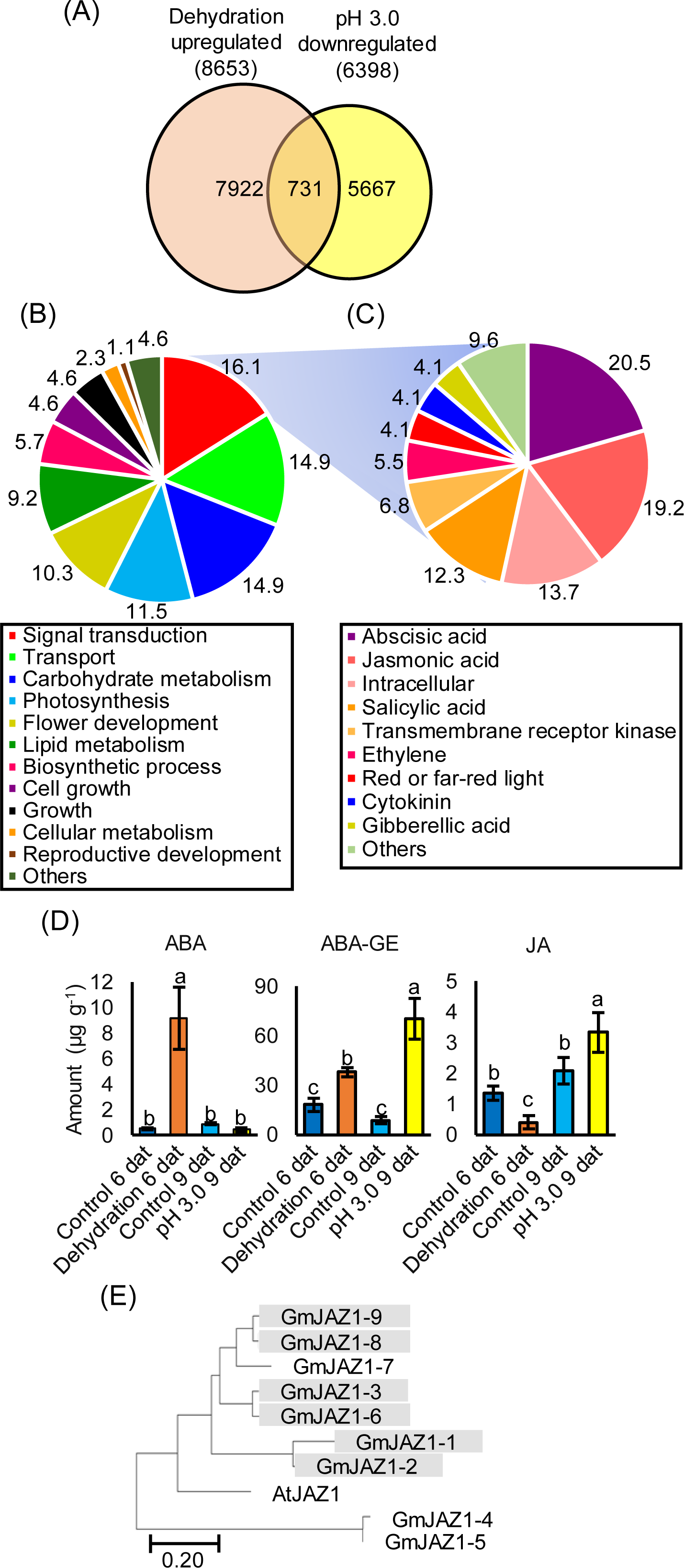
Comparative transcriptomics of soybean seedlings treated with dehydration for 6 d or acidity stress (pH 3.0 medium) for 9 d. (A) Venn diagram of genes from Harosoy 63 seedlings that were upregulated by dehydration and/or downregulated by acidity stress. (B) Pie charts showing the percentage of genes by ontology that were up- and down-regulated by the stresses. (C) A further breakdown of the ‘Signal Transduction’ ontology into subcategories. (D) ABA, ABA-GE, and JA metabolite amounts in Harosoy 63 seedlings after 6 d dehydration and 9 d acidy stress measured by UPLC-MS^n^. Statistical analysis was run separately for each hormone. Different letters show significant differences by single factor ANOVA, Tukey post hoc test, *P* < 0.05, α = 0.05. Error bars = standard error (SE), n = 3-5 per condition. A repeat of this experiment provided similar results and is shown in Supplementary Figure S1. For a list of statistical values for both experiments, see Supplementary Table S1. (E) Cluster analysis of deduced amino sequences of GmJAZ1s with Arabidopsis AtJAZ1. Branches represent estimated evolutionary relationships based on amino acid sequence similarity. The scale bar indicates the number of differences per 100 residues derived from the Muscle alignment. Gray boxes represent GmJAZ1s that were up- and down-regulated in Harosoy 63 seedlings by 6 d dehydration and 9 d acidity stresses, respectively.

**Table 1.**
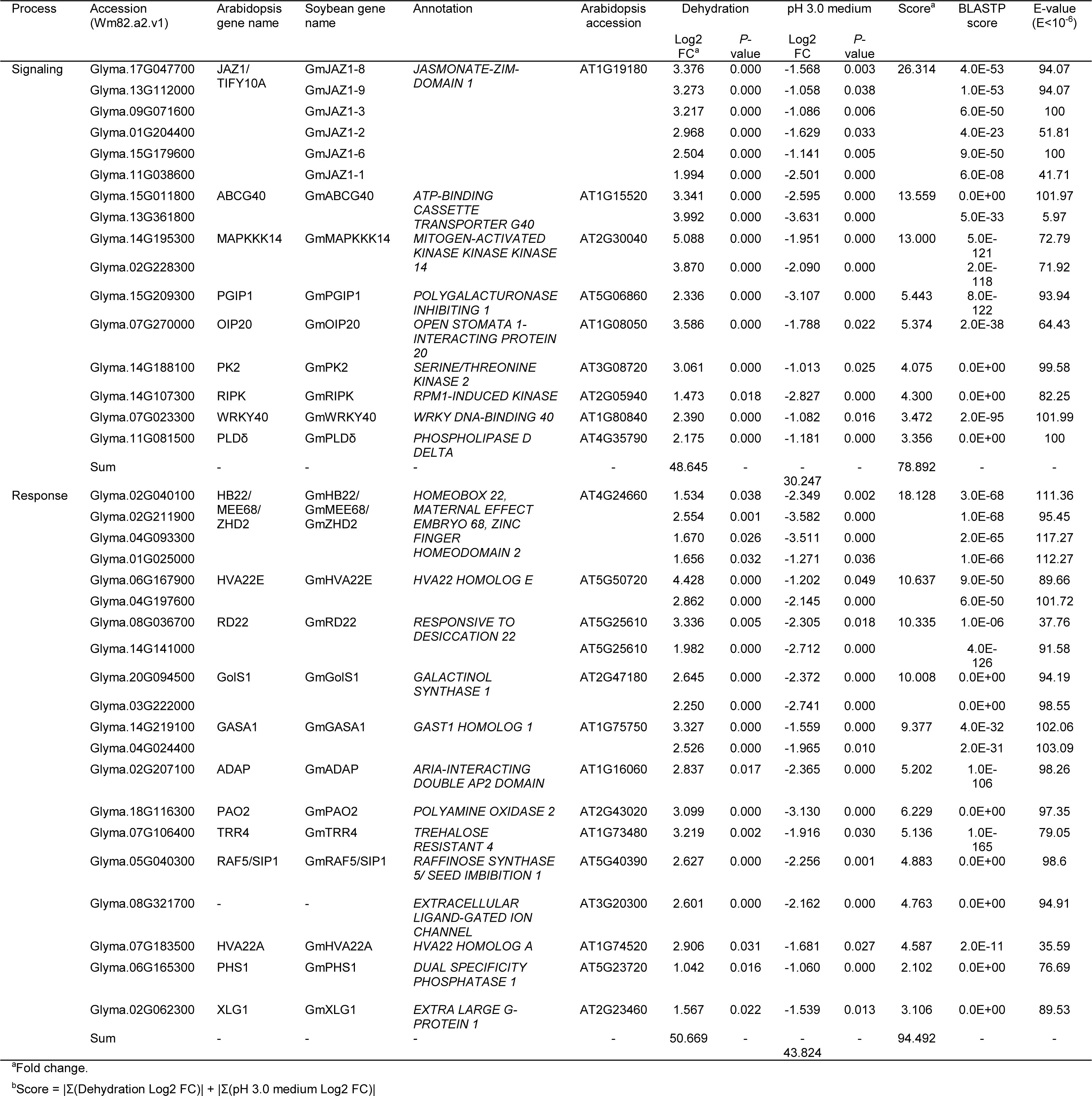
GmJAZ1 and ABA genes upregulated by dehydration and downregulated by pH 3.0 medium.

Many JA marker genes demonstrated expression patterns that were similar to JA metabolite accumulation (Table 2, Figure 2D), and glyceollin biosynthesis (Jahan *et al*., 2019). This is consistent with the role of JA in positively regulating phytoalexin biosynthesis (Zhou et al., 2022). However, since JAZ1 genes negatively regulate phytoalexin biosynthesis in Arabidopsis and grapevine, this drew our attention to the six GmJAZ1 genes that did not follow JA marker gene expressions, but rather had expression profiles similar to ABA genes and metabolites (Table 2, Figure 2D). To understand the relatedness among the predicted GmJAZ1 proteins and AtJAZ1 from Arabidopsis, we conducted a cluster analysis based on their amino acid sequences. Seven out of nine GmJAZ1s formed a subgroup with AtJAZ1 (Figure 2E). All six *GmJAZ1* genes that were upregulated by dehydration stress encoded proteins that clustered in a subgroup with AtJAZ1. Thus, we considered these six *GmJAZ1* genes and ABA signaling as candidate negative regulators of glyceollin biosynthesis.

**Table 2.**
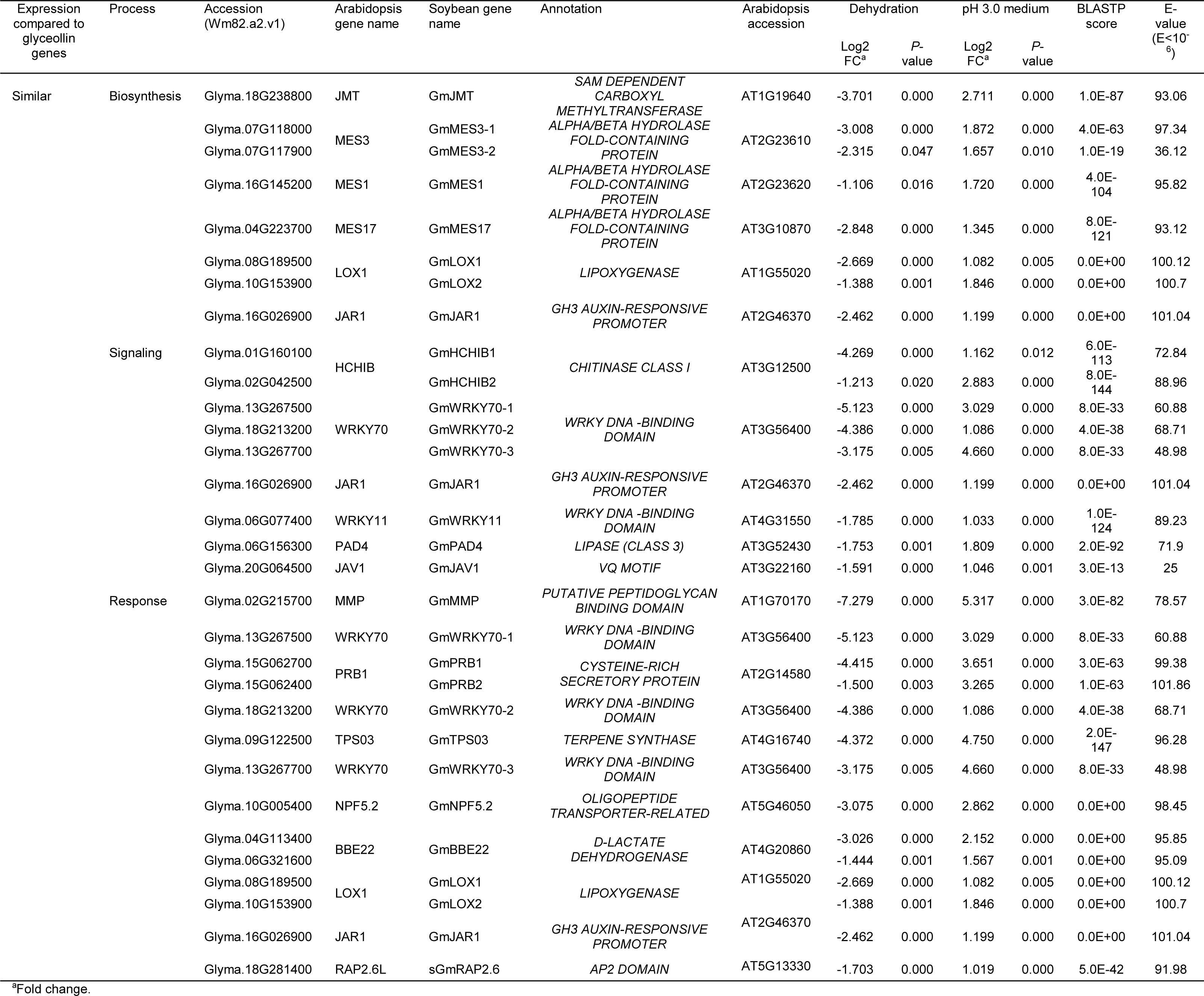
JA genes downregulated by dehydration and upregulated pH 3.0 medium.

### Suppression of glyceollin biosynthesis by dehydration stress is dependent on ABA and ABA-upregulated *GmJAZ1* genes

To confirm whether ABA suppresses the elicitation of glyceollin biosynthesis, we transferred soybean seedlings to pH 3.0 elicitation medium with and without ABA, and grew the seedlings for 6 d. The inclusion of ABA in the medium reduced glyceollin metabolite amounts 5.8- to 12.4-fold, resulting in similar levels compared to the non-eliciting pH 5.8 control medium (Figure 3A). Statistical values for these experiments are shown in Supplementary Table S2. ABA also suppressed the accumulation of the phytoalexins β-prenyl-genistein and phaseol, and the constitutively accumulating isoflavonoids 6“- *O*-malonylononin and 6”-*O*-malonyldaidzin (Figure 3A). It did not suppress 6“-O-malonylgenistin and increased the accumulation of an unknown metabolite. This metabolite exhibited UV absorbance properties similar to isoflavonoids, but did not represent any of the 57 (iso)flavonoid standards that we have in our library. To test whether ABA could suppress the elicitation of glyceollins in response to a biotic elicitor, we treated soybean hairy roots with *P. sojae* WGE with and without ABA. The ABA treatment suppressed WGE’s elicitation of total glyceollins 16.1-fold (Figure 3B). A repeat of this experiment demonstrated similar results (Supplementary Figure S2). Statistical values for these experiments are shown in Supplementary Table S3. Thus, exogenous ABA treatment suppresses both abiotic and biotic elicitation of glyceollins.

**Figure 3.**
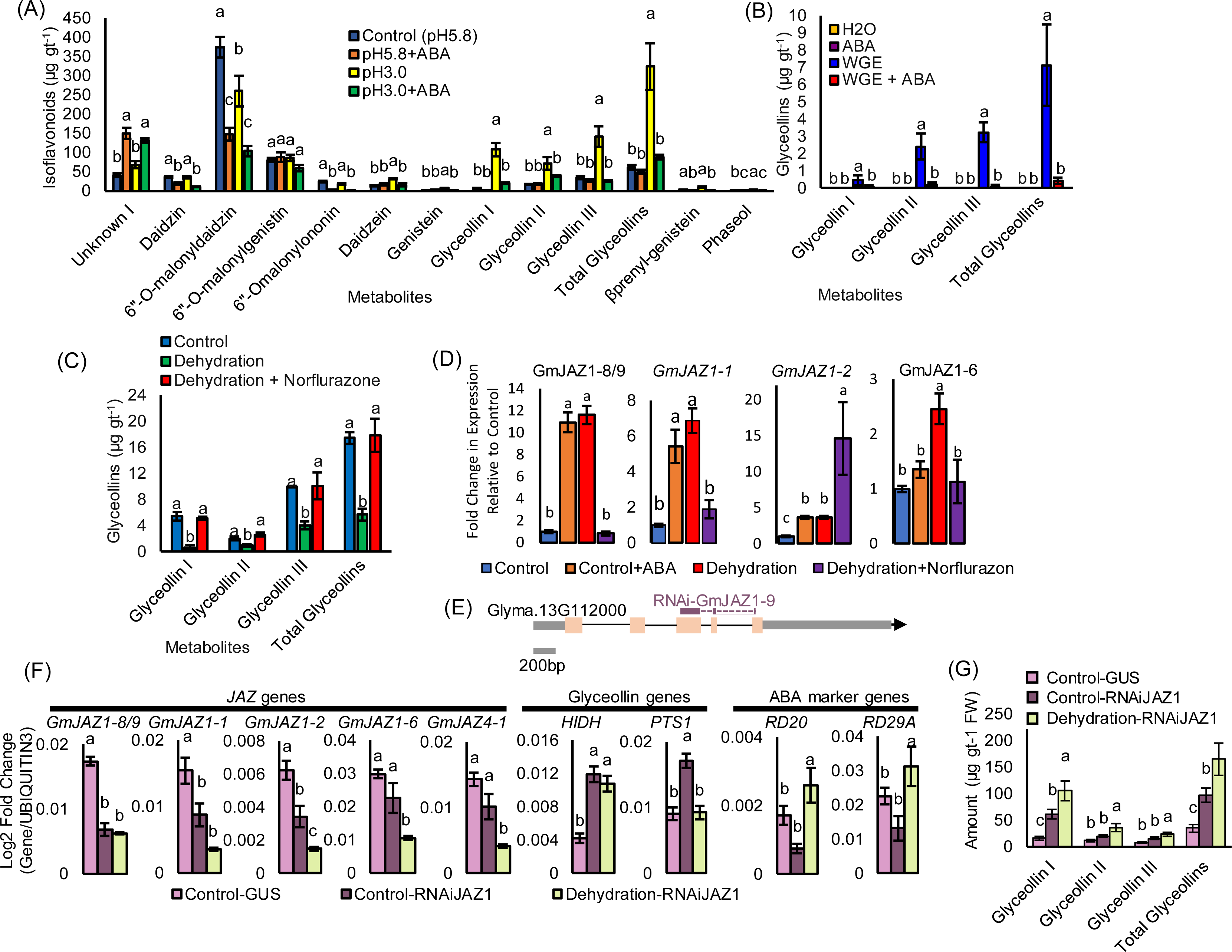
ABA and ABA-upregulated *GmJAZ1* genes suppress the elicitation of glyceollin biosynthesis. (A) Isoflavonoid metabolite amounts in Harosoy 63 seedlings treated for 9 d with pH 3.0 medium with and without ABA. Different letters show significant differences by single factor ANOVA, Tukey post hoc test, P < 0.05, α = 0.05. Error bars = standard error (SE), n ≥ 3 per condition. For a list of statistical values for this experiment, see Supplementary Table S2. (B) Glyceollin amounts in W82 soybean hairy roots after 24 h treatment with *Phytophrhora sojae* wall glucan elicitor (WGE) with and without ABA. Different letters show significant differences by single factor ANOVA, Tukey post hoc test, P < 0.05, α = 0.05. Error bars = standard error (SE), n ≥ 3 per condition. A repeat of this experiment is shown in Supplementary Figures S2. For a list of statistical values from both experiments, see Supplementary Table S3. (C) Glyceollin metabolite amounts in Harosoy 63 soybean seedlings after 6 d dehydration with or without norflurazone treatment measured by UPLC-PDA. Different letters show significant differences by single factor ANOVA, Tukey post hoc test, *P* < 0.05, α = 0.05. Error bars = standard error (SE), n ≥ 3 per condition. A repeat of this experiment is shown in Supplementary Figures S3. For a list of statistical values for this experiment, see Supplementary Table S4. (D) Expression levels of *GmJAZ1* genes in W82 seedlings upon exposure to 6 d dehydration, dehydration+norflurazon, or ABA treatment measured by qRT-PCR. Different letters show significant differences by single factor ANOVA, Tukey post hoc test, *P* < 0.05, α = 0.05. Error bars represent SE (n = 5 per condition). For statistical values see Supplementary Table S5. (E) Schematic diagram of the *GmJAZ1-9* gene and the region used go clone the 302 bp RNAi trigger site. (F) Expression levels of GmJAZ1 genes and glyceollin biosynthesis genes in transgenic W82 soybean seedlings after 6 days of dehydration treatment, measured by qRT-PCR. Different letters show significant differences by single factor ANOVA, Tukey post hoc test, *P* < 0.05, α = 0.05. Error bars represent SE (n = 5 per condition). For a list of statistical values from both experiments, see Supplementary Table S7. (G) Glyceollin levels in transgenic W82 soybean hairy roots transformed with an empty vector or an RNA interference construct targeting the JAZ1 gene, measured after 6 days of treatment using UPLC-PDA. Different letters show significant differences by single factor ANOVA, Tukey post hoc test, *P* < 0.05, α = 0.05. Error bars = standard error (SE), n ≥ 3 per condition. For a list of statistical values for this experiment, see Supplementary Table S8.

To determine whether dehydration stress employs ABA signaling to suppress glyceollin biosynthesis, we treated soybean seedlings with the ABA biosynthesis inhibitor norflurazon. Dehydration stress reduced the amounts of total glyceollins by 3.1-fold, whereas the norflurazon pretreatment completely prevented the suppressive effects of dehydration (Figure 3C). A repeat of this experiment demonstrated similar results (Supplementary Figure S3). Statistical values for these experiments are shown in Supplementary Table S4. These results demonstrate that the suppression of glyceollin biosynthesis by dehydration stress is dependent on ABA biosynthesis. Since ABA and *GmJAZ1* genes were upregulated by dehydration stress in our RNA-seq experiments, we tested whether dehydration stress and ABA upregulate the expression *GmJAZ1* genes. ABA treatment for 6 d, which is the same duration of dehydration treatment that was used for our RNA-seq study, increased the expression of *GmJAZ1-8/9* and *GmJAZ1-1* by 10.9- and 4.5-fold, respectively (Figure 3D). For statistical results see Supplementary Table S5. Notably, after 24 h of ABA treatment, which is the time point typically used to study the effects of ABA treatments, the expression of *GmJAZ1* transcripts did not change, despite that some other ABA-responsive genes were upregulated (Supplementary Figure S4). By contrast, exposing soybean seedlings to dehydration stress for 6 d upregulated *GmJAZ1-8/9*, *GmJAZ1-1*, *GmJAZ1-2*, and *GmJAZ1-6* 2.5- to 11.6-fold, with *GmJAZ1-8/9* being the most highly upregulated. To test whether the upregulation is ABA-dependent, we pretreated seedlings with norflurazon and subjected them to dehydration stress. Adding norflurazon to the seedling growth medium prior to dehydration treatment completely abolished the increase in expression of *GmJAZ1-8/9*, *GmJAZ1-1*, and *GmJAZ1-6* (Figure 3D). These results indicate that dehydration stress employs ABA signaling to upregulate the expression of *GmJAZ1-8/9*, *GmJAZ1-1*, and *GmJAZ1-6*. By contrast, *GmJAZ1-2* is upregulated by norflurazon, potentially suggesting functional divergence among some homologs.

Since dehydration stress and ABA treatment upregulated several *GmJAZ1* genes, we tested whether the suppression of glyceollin biosynthesis by dehydration is dependent on *GmJAZ1s*. We made soybean composite plants that consist of non-genetically modified aerial tissues attached to genetically modified hairy roots that were transformed with an *RNAi-GmJAZ1* gene silencing construct. We chose to clone a 302 bp fragment of the coding sequence of *GmJAZ1-9* to use as the RNAi trigger for the *RNAi-GmJAZ1* construct (Figure 5E). This is because *GmJAZ1-8/9* were most highly upregulated by dehydration and ABA, and their upregulation by dehydration was dependent on ABA. Since the cleavage of mRNA targets by RNAi depends on 21-nucleotide base pairings among the RNAi trigger and the target gene, we calculated the number of non-overlapping 21-nucleotide pairings that span the potential *GmJAZ1* target genes. *GmJAZ1-9* and *GmJAZ1-8* had 14 and nine predicted 21-nucleotide stretches that were complementary to the *RNAi-GmJAZ1* trigger (Supplementary Table S6). By contrast, only one pairing was observed for each of the other *GmJAZ1s*, and the off-target control *GmJAZ4-1* had zero pairings. Based on this, we predicted that our *RNAi-GmJAZ1* construct would target most efficiently *GmJAZ1-9* and *GmJAZ1-8*. The trigger was cloned into the RNAi vector *pANDA35HK* in inverted orientations to generate an RNAi hairpin when expressed, and the resultant *RNAi-GmJAZ1* construct was transferred into soybean to make composite plants.

**Figure 4.**
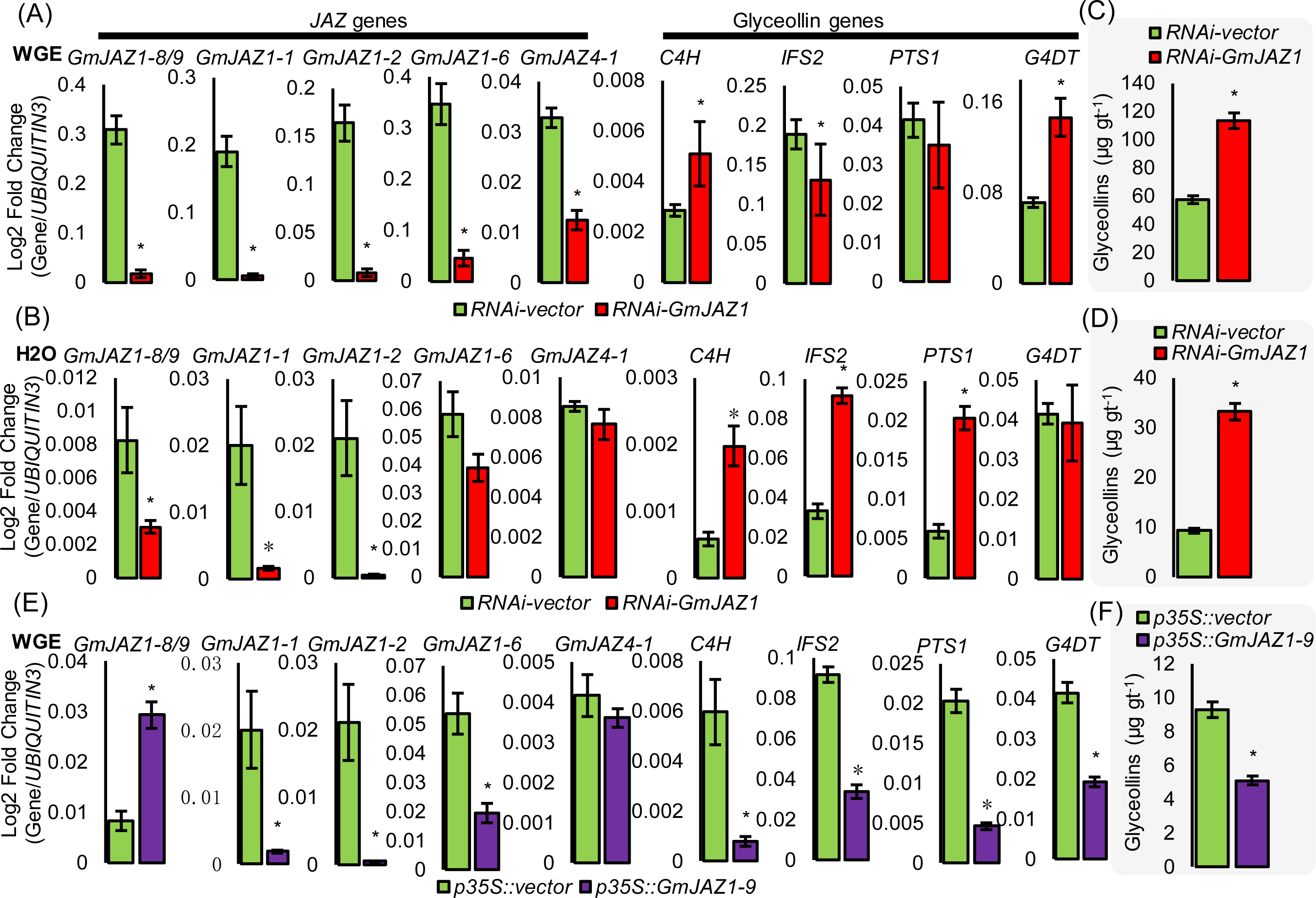
GmJAZ1s are negative regulators of glyceollin phytoalexin biosynthesis in soybean. (A) Gene expressions in *RNAi-GmJAZ1* W82 hairy roots elicited for 24 h with WGE. Results represent the average expression of five independent hairy root transformation events. *Significantly different than control by paired Student’s *t*-test (P < 0.05). Error bars represent SE (n = 5 per genotype). A repeat of this experiment is shown in Supplementary Figure S5. For a list of statistical values from both experiments, see Supplementary Table S9. (B) Gene expressions in 24 h mock (H2O)-treated RNAi-GmJAZ1 W82 hairy roots. Results represent the average expression of five independent hairy root transformation events. *Significantly different than control by paired Student’s *t*-test (P < 0.05). Error bars represent SE (n = 5 per genotype). For a list of statistical values, see Supplementary Table S10. (C) Amount of total glyceollin metabolites in RNAi-GmJAZ1 W82 hairy roots elicited for 24 h with WGE measured by UPLC-PDA. Results represent the average expression of five independent hairy root transformation events. *Significantly different than control by paired Student’s *t*-test (P < 0.05). Error bars represent SE (n = 5 per genotype). A repeat of this experiment is shown in Supplementary Figures S5. For a list of statistical values from both experiments, see Supplementary Table S11. (D) Amounts of total glyceollin metabolites in 24 h mock-treated RNAi-GmJAZ1 W82 hairy roots. *Significantly different than control by paired Student’s *t*-test (P < 0.05). Error bars represent SE (n = 5 per genotype). For a list of statistical values, see Supplementary Table S12. (E) Gene expressions in W82 hairy roots overexpressing GmJAZ1-9 elicited for 24 h with WGE. Results represent the average expression of five independent hairy root transformation events. *Significantly different than control by paired Student’s *t*-test (P < 0.05). Error bars represent SE (n = 5 per genotype). A repeat of this experiment is shown in Supplementary Figures S5. For a list of statistical values from both experiments, see Supplementary Table S13. (F) Total glyceollin amounts in W82 hairy roots overexpressing GmJAZ1-9 after 24 h WGE treatment. *Significantly different than control by paired Student’s *t*-test (P < 0.05). Error bars represent SE (n = 5 per genotype). A repeat of this experiment is shown in Supplementary Figures S5. For a list of statistical values from both experiments, see Supplementary Table S14.

**Figure 5.**
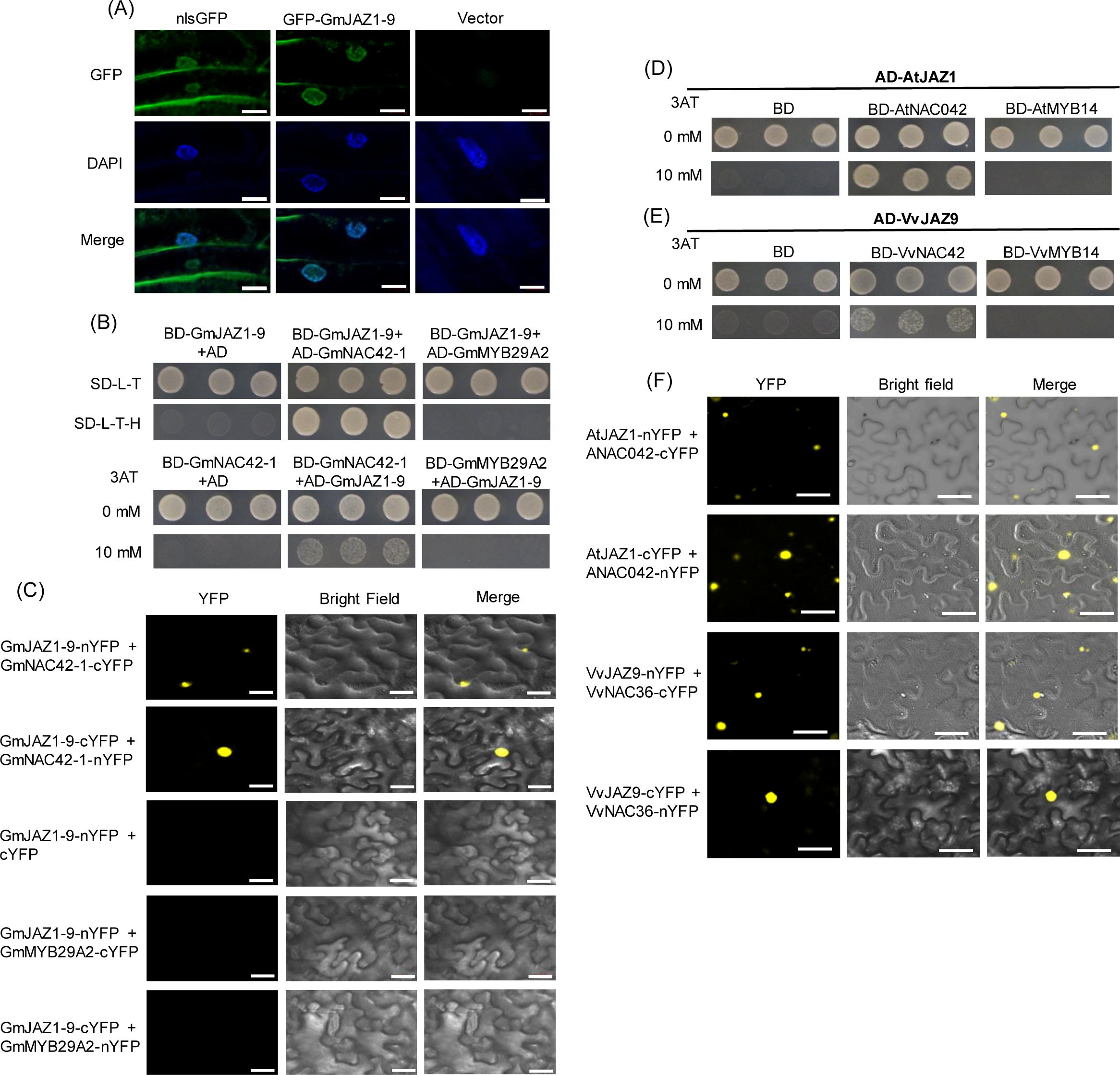
JAZ1 proteins from soybean, Arabidopsis and grapevine interact with NAC42 protein activators of phytoalexin biosynthesis. (A) Fluorescence microscopy of GmJAZ1-9 translationally fused to an N-terminal green fluorescent protein (GFP) tag in transgenic W82 hairy roots. A nuclear localization signal translationally fused to GFP (NLS-GFP) was used as positive control. An empty vector was used as negative control. DAPI (6 µg/ml) demonstrates nuclear staining. (B) Yeast two-hybrid analysis testing for interactions among GmJAZ1-9 and GmNAC42-1 or GmMYB29A2 proteins. Yeast strain PJ69-4a was co-transformed with genes that were translationally fused to the GAL4 activation domain (AD) or binding domain (BD). Yeast containing plasmid pairs were plated on SD-Leu-Trp (SD-L-T) and SD-Leu-Trp-His (SD-L-T-H) ±3-aminotriazole (3AT). Growth on SD-L-T or SD-L-T-H+ 0 mM 3AT (top rows) are plating controls, on SD-L-T-H or SD-L-T+H + 10 mM 3AT (bottom rows) indicates positive protein-protein interactions. pDEST-GADT7 (AD) was used as an empty ‘vector’ negative control. 3AT is used for attenuating auto-activation. (C) BIFC assays in *N. benthamiana* to confirm the interactions of GmJAZ1-9 and GmNAC42-1. (D) Yeast two-hybrid analysis testing for interactions between Arabidopsis AtJAZ1 and ANAC042 or AtMYB14. Yeast strain PJ69-4a was co-transformed and plated on SD-L-T-H and SD-L-T-H ±3AT. Yeast two-hybrid analysis testing for interactions between VvJAZ9 and VvNAC42 or VvMYB14. Yeast strain PJ69-4a was transformed and plated on SD-L-T-H and SD-L-T-H ±3AT. Growth on SD-L-T-H+ 0 mM 3AT (top rows) are plating controls, on SD-L-T+H + 10 mM 3AT (bottom rows) indicates positive protein-protein interactions. **F**, BIFC assays to confirm the interactions of AtJAZ1 and ANAC042, and VvJAZ9 and VvNAC42. All images were collected using Zeiss confocal microscope. Scale bars are 20 µm. Additional images and negative controls for C and F are shown in Supplementary Figure S6.

The hairy roots of composite plants transformed with *RNAi-GmJAZ1* exhibited a 1.76- to 2.54-fold reduction in the expression of *GmJAZ1-8/9*, *GmJAZ1-1*, and *GmJAZ1-2,* with GmJAZ1-8/9 showing the greatest reduction, and no change in expression of *GmJAZ1-6* and *GmJAZ4-1* (Figure 3F). By contrast, they exhibited a 2.85- and 1.89-fold increase in expression of the glyceollin biosynthetic genes *HIDH* and *PTS1* (Figure 3F), and a 4.7-fold increase in the amounts of total glyceollins (Figure 3G). For statistical results see Supplementary Tables S7 and S8. Finally, dehydration treatment failed to suppress glyceollin biosynthesis in *RNAi-GmJAZ1* composite plants (Figures 3F, 3G). ABA marker genes were significantly upregulated in the dehydration treated plants, indicating the dehydration treatment stimulated ABA signaling. These results demonstrate that the suppression of glyceollin biosynthesis by dehydration stress is dependent on the expression of *JAZ1* genes *GmJAZ1-8/9*, *GmJAZ1-1*, and/or *GmJAZ1-2*.

### Silencing *GmJAZ1-8/9/1/2* derepresses and overexpressing *GmJAZ1-9* suppresses glyceollin biosynthesis

To further investigate whether *GmJAZ1* genes have a role in regulating glyceollin biosynthesis, we generated soybean hairy roots transformed with the *RNAi-GmJAZ1* construct and treated them with *P. sojae* WGE. The soybean hairy root response to WGE has served extensively as a model system for investigating the genetic regulation of glyceollin biosynthesis (Graham *et al*., 2007, Jahan *et al*., 2019, Jahan *et al*., 2020, Lin *et al*., 2023a, Lin *et al*., 2024). Under WGE treatment, *RNAi-GmJAZ1* hairy roots exhibited a 17.57- to 29.71-fold reduction in the expression of *GmJAZ1-8/9*, *GmJAZ1-1*, and *GmJAZ1-2* (Figure 4A). They also exhibited a 7.68- and 2.54-fold reduction in the expression of *GmJAZ1-6* and *GmJAZ4-1* (Figure 4A). Note, *GmJAZ1-6* and *GmJAZ4-1* did not have reduced expression in mock-treated *RNAi-GmJAZ1* hairy roots (Figure 4B) or *RNAi-GmJAZ1* composite plants (Figure 3F), suggesting that their reduced expression may be due to effects of WGE on downstream signaling regulated by direct targets of *RNAi-GmJAZ1*. The *RNAi-GmJAZ1* hairy roots treated with WGE had a 1.8- and 2.1-fold increase in the expressions of the glyceollin biosynthesis genes *C4H* and *G4DT* and a 2.0-fold increase in the amounts of total glyceollin metabolites (Figure 4A, 4C, for statistical results see Supplementary Table S9 and S11). A repeated experiment demonstrated similar results (Supplementary Figure S5, Supplementary Table S9 and S11).

Under mock treatment, the *RNAi-GmJAZ1* hairy roots exhibited a 2.75- to 53.43-fold reduction in the expression of *GmJAZ1-8/9*, *GmJAZ1-1*, and *GmJAZ1-2* (Figure 4B). This was accompanied by a 2.76- to 3.4-fold upregulation of the glyceollin biosynthetic genes *C4H*, *IFS2*, and *PTS1* and a 3.6-fold increase in glyceollin metabolite amounts (Figures 4B, 4D, for statistical results see Supplementary Tables S10 and S12). Notably, glyceollin levels in the mock-treated *RNAi-GmJAZ1* roots were 60% of the WGE-elicited control (Figures 4D). These results demonstrate that silencing *GmJAZ1-8/9*, *GmJAZ1-1*, and *GmJAZ1-2* derepresses glyceollin biosynthesis in non-elicited tissues.

To investigate further whether *GmJAZ1s* can negatively regulate glyceollin biosynthesis, we overexpressed the coding sequence of *GmJAZ1-9* in soybean hairy roots using the viral *CaMV 35S* promoter. Overexpressing *GmJAZ1-9* 2.7-fold in elicited roots resulted in a 2.2- to 7.6-fold reduction in transcript levels of glyceollin biosynthesis genes and a 3.4-fold reduction in glyceollin metabolites (Figure 4E, 4F, for statistical results see Supplementary Tables S13 and S14). Also, it reduced the expressions of *GmJAZ1-1*, *GmJAZ1-2*, and *GmJAZ1-6* (Figure 4E), demonstrating that the ectopic overexpression of *GmJAZ1-9* alone is sufficient to suppress glyceollin biosynthesis. While further experiments would be needed to dissect the relative functions of each *GmJAZ1-8, GmJAZ1-1, and GmJAZ1-2*, our overexpression results strongly support that *GmJAZ1-9* has a role in negatively regulating glyceollin biosynthesis.

### The JAZ1-NAC42 protein-protein interaction is conserved among plant species and suppresses the activation of glyceollin biosynthesis in soybean

Previous studies have found that JAZ proteins physically interact with transcriptional activators (*e.g.* MYB and bHLH proteins) and inhibit their transactivation activities (Fernández-Calvo *et al*., 2011, Niu *et al*., 2011, Qi *et al*., 2015, Zhang *et al*., 2015, Li *et al*., 2021). Since silencing *JAZ1* genes derepresses phytoalexin biosynthesis in Arabidopsis (Makhazen *et al*., 2021a), grapevine (Makhazen *et al*., 2021b), and soybean (Figures 3, 4), we hypothesized that GmJAZ1 proteins function by physically interacting with one or more conserved transcriptional activators of phytoalexin biosynthesis. We first determined whether the subcellular localization of GmJAZ1-9 is consistent with this role. We assembled a translational fusion of the GmJAZ1-9 CDS with an N-terminal GFP tag and investigated its subcellular localization by confocal microscopy. GFP-GmJAZ1-9 localized to the nucleus of soybean hairy root cells (Figure 5A). To determine whether the GmJAZ1-9 protein physically interacts with known transcriptional activators of glyceollin biosynthesis, we conducted yeast two-hybrid (Y2H) experiments with GmNAC42-1 and GmMYB29A2. Each CDS was translationally fused with N-terminal tags of GAL4 activation domain (AD) and binding domain (BD). Using this approach, GmJAZ1-9 physically interacted with GmNAC42-1, but not with GmMYB29A2 (Figure 5B). The interactions were observed when GmJAZ1-9 was fused to either GAL4 AD or BD. The same interactions were observed in the nucleus by bimolecular fluorescence complementation (BiFC) assays when the proteins were expressed as translational fusions with C-terminal split yellow fluorescent protein (YFP) tags in the *Nicotiana benthamiana* system (Figure 5C, Supplementary Figure S6). These results show that GmJAZ1-9 and GmNAC42-1 physically interact in the nucleus. We also tested whether the JAZ1 proteins from Arabidopsis and grapevine interact with their corresponding NAC42 transcription factors. Arabidopsis ANAC042 interacted with JAZ1, and grapevine VvNAC36 interacted with VvJAZ9 in both the Y2H and *N. benthamiana* BiFC systems (Figure 5D-F, Supplementary Figure S6). Together, our results demonstrate that the JAZ1-NAC42 interaction is conserved among soybean, Arabidopsis, and grapevine proteins.

Since JAZ proteins have been found to suppress the transactivation activities of several transcription factors (Fernández-Calvo *et al*., 2011, Niu *et al*., 2011, Qi *et al*., 2015, Zhang *et al*., 2015, Li *et al*., 2021), we tested whether GmJAZ1-9 suppresses GmNAC42-1’s transactivation of glyceollin gene promoters. We first inserted the promoters of the glyceollin biosynthetic genes *2−HYDROXY-ISOFLAVANONE DEHYDRATASE* (*pHIDH*) and *PTEROCARPAN SYNTHASE1* (*pPTS1*) into a vector upstream of the luciferase reporter gene. We then co-transfected the DNA constructs into cucumber protoplasts with plasmids expressing *GmNAC42-1* alone or with *GmJAZ1-9*. Co-transfection of *GmNAC42-1* with the empty vector found that *GmNAC42-1* activated both *pHIDH* and *pPTS1* promoters (Figure 6A). Importantly, co-transfection with *GmJAZ1-9* reduced GmNAC42-1’s transactivation of both promoters by 83.5 – 89.0% compared to the *GmNAC42-1*/empty vector control (Figure 6A). GmMYB29A2 activated *pPTS1* but not *pHIDH* (Figure 6B). In contrast to GmJAZ1-9’s effect on GmNAC42-1 activity, the co-transfection of *GmJAZ1-9* with *GmMYB29A2* did not affect GmMYB29A2’s transactivation activity (Figure 6B). This is consistent with the finding that GmJAZ1-9 and GmMYB29A2 proteins do not interact in Y2H and BiFC experiments (Figure 5B, 5C). To determine whether GmJAZ1-9 specifically affected the DNA binding ability of GmNAC42-1, we conducted yeast three-hybrid (Y3H) assays. Co-transforming *GmJAZ1-9* with *GmNAC42-1* reduced GmNAC42-1’s binding to the *pHIDH* and *pPTS1* promoters compared to the GmNAC42-1/empty vector control (Figure 6C). However, GmJAZ1-9 did not affect GmMYB29A2’s binding to those promoters (Figure 6C). Collectively, the results show that nuclear-localized GmJAZ1-9 suppresses GmNAC42-1’s DNA binding and transactivation activities to suppress the activation of glyceollin biosynthetic gene promoters.

**Figure 6.**
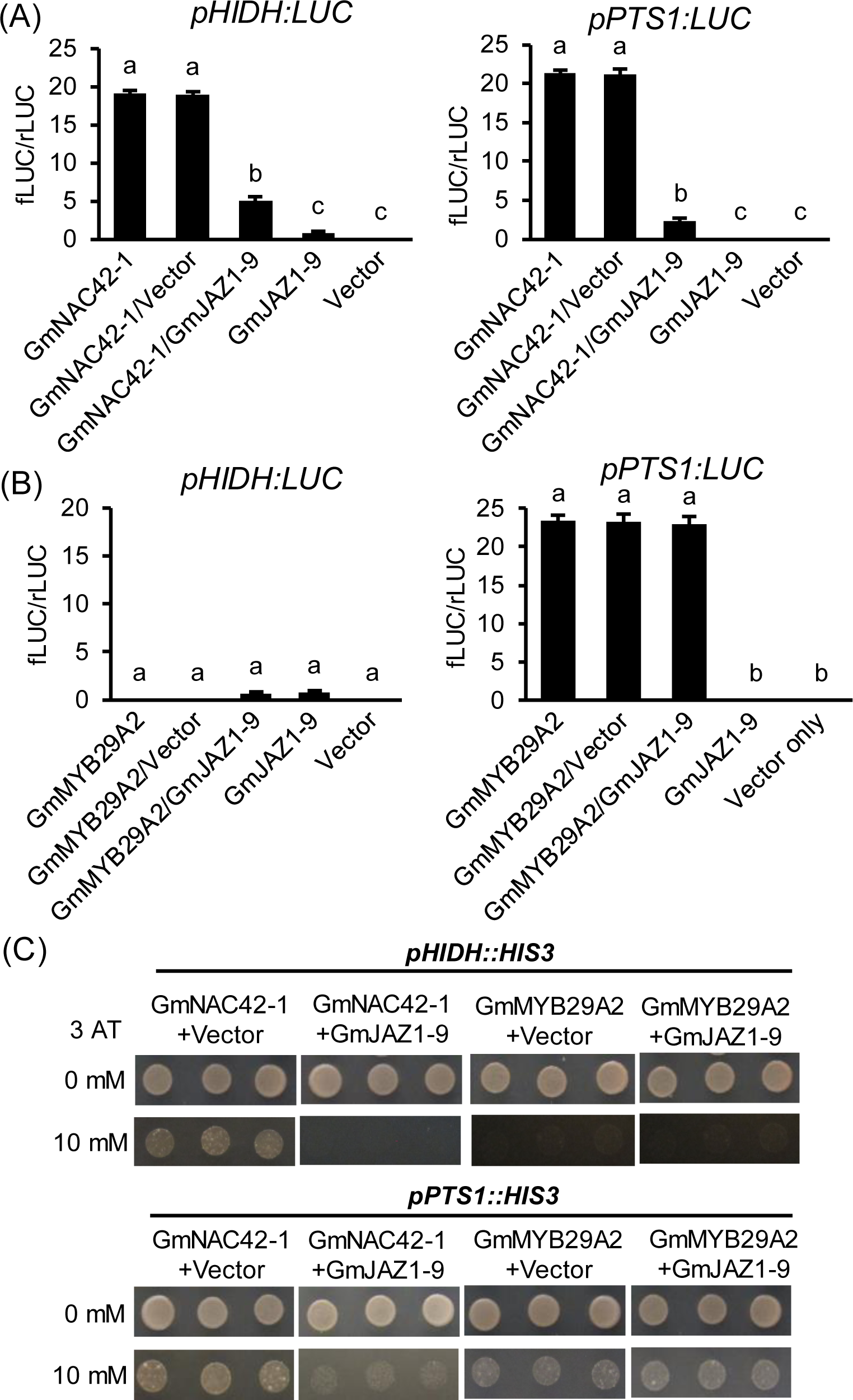
GmJAZ1-9 inhibits GmNAC42-1 transactivation of glyceollin biosynthesis gene promoters by inhibiting its DNA binding. (A) Effects of GmJAZ1-9 on GmNAC42-1’s transactivation of pHIDH and pPTS1 promoters in transfected cucumber protoplasts. Values on the y axis are normalized to co-transfected Renilla luciferase reporter construct. The error bars designate SE (n = 4 per construct). Different letters show significant differences by single factor ANOVA, Tukey post hoc test, *P* < 0.05, α = 0.05. All promoters comprise the 500-bp region upstream of their respective translational start sites. (B) Effects of GmJAZ1-9 on GmMYB29A2 transactivation of pHIDH and pPTS1 promoters in transfected cucumber protoplasts. The error bars designate SE (n = 4 per construct). Different letters show significant differences by single factor ANOVA, Tukey post hoc test, *P* < 0.05, α = 0.05. (C) Yeast three-hybrid analysis of strain PJ69-4a transformed with GmNAC42-1-Gal4AD and GmJAZ1-9 on SD-Leu-His-Ura (SD-L-H-U) and SD-L-H-U ±3-aminotriazole (3AT). Growth on SD-L-H-U + 10 mM 3AT (top rows) are plating controls, and SD-L-H-U + 10 mM 3AT (bottom rows) indicate positive protein-DNA interactions. pDR-GW was used as the empty ‘Vector’ control.

## DISCUSSION

Decades ago, dehydration stress, and its major signaling hormone, ABA, were found to be negative regulators of different phytoalexin biosynthetic pathways in diverse plant lineages (Henfling *et al*., 1981, Goossens *et al*., 1987, Mohr and Cahill, 2003, Asselbergh *et al*., 2007, de Torres Zabala *et al*., 2009, Fan *et al*., 2009). Here, our RNA-seq analysis of soybean seedlings comparing the effects of long-term (6 d) dehydration stress and elicitor treatment found that ABA signaling genes are oppositely expressed compared to glyceollin biosynthetic genes (Figure 2B, 2C, Table 1). The results implicated ABA signaling in negatively regulating the biosynthesis of glyceollins, soybean’s major phytoalexins. In support of this role, our results demonstrated that pretreating soybean seedlings with an ABA biosynthesis inhibitor prevented dehydration stress from suppressing glyceollin biosynthesis (Figure 3C). This supported that the suppression of glyceollin biosynthesis by dehydration is dependent on endogenous ABA biosynthesis. Further support of this suppressive role came from ABA treatments that suppressed glyceollin biosynthesis in soybean seedlings that were elicited with acidity stress and soybean hairy roots elicited with *P. sojae* wall glucan elicitor (WGE) (Figure 3A, 3B). Thus, ABA treatment can suppress both abiotic and biotic elicitation of glyceollin biosynthesis. A role of ABA in negatively regulating glyceollin biosynthesis may be relevant to soybean resistance in agriculture. Glyceollins have a major role in conferring resistance to *P. sojae* and ABA negatively regulates soybean’s resistance (Mohr *et al*., 2001, Graham *et al*., 2007, Jahan *et al*., 2020). Further, active levels of ABA are depleted in incompatible soybean genotypes during the response to *P. sojae* (McDonald and Cahill, 1999, Mohr *et al*., 2001, Asselbergh *et al*., 2007). Thus, our findings suggest that dehydration stress suppresses glyceollin biosynthesis by enhancing ABA signaling that is typically active prior to elicitation. They also raise the possibility that long-term dehydration stress, and potentially other stresses that are communicated by ABA, such as salt and cold stresses, may predispose soybean plants to subsequent infection by *P. sojae*. To our knowledge this has not yet been investigated. Some pathogens exploit ABA biosynthesis to promote compatibility. The AvrPtoB effector secreted from *P. syringae* stimulates the expression of ABA biosynthesis to promote compatibility (de Torres-Zabala *et al*., 2007). Thus, conditional silencing of ABA genes, potentially those identified by our RNA-seq experiments, could be valid candidates for crop improvement by breeding or bioengineering efforts.

Our RNA-seq analysis of soybean seedlings comparing the effects of long-term dehydration stress and elicitor treatments found that *GmJAZ1* genes are oppositely expressed compared to glyceollin biosynthetic genes (Figure 2B, 2C, Table 1). The results implicated *GmJAZ1* genes in negatively regulating glyceollin biosynthesis. RNAi silencing of *JAZ1* genes in soybean hairy roots demonstrated that the suppression of glyceollin biosynthesis by dehydration stress is dependent on *GmJAZ1-8/9*, *GmJAZ1-1*, and/or *GmJAZ1-2* (Figure 3F, 3G). More insight into the role of *GmJAZ1* genes came from experiments that lacked an elicitor treatment. Both soybean composite plants and hairy roots grown *in vitro* demonstrated that silencing *GmJAZ1-8/9*, *GmJAZ1-1*, and *GmJAZ1-2* in control conditions, without and elicitor treatment, derepresses glyceollin biosynthesis (Figure 3F, 3G, 4B, 4D). Similar findings were recently reported for Arabidopsis and grapevine. RNAi silencing of *JAZ1* genes in Arabidopsis calli and grapevine cell suspensions resulted in the derepression of camalexin and stilbene phytoalexin biosynthesis, respectively (Makhazen *et al*., 2021a, Makhazen *et al*., 2021b). Taken together, these studies and our results in soybean (Figure 3F, 3G, 4B, 4D), suggest that *JAZ1* genes have a conserved role in keeping phytoalexin biosynthesis ‘off’, or at basal levels, in the absence of elicitation. Further, our results support an additional role of *GmJAZ1s*, specifically in elicited soybean tissues. Our RNAi experiments demonstrate that, following elicitation, silencing *GmJAZ1-8/9, GmJAZ1-1, and GmJAZ1-2* results in increased levels of glyceollin biosynthesis (Figure 4A, 4C). Thus, in addition to keeping glyceollin synthesis ‘off’ in the absence of elicitation, GmJAZ1s function to limit the level of glyceollin biosynthesis following elicitation. These results distinguish the roles of GmJAZ1s from those of GmWRKY72, that limits glyceollin biosynthesis only upon elicitation (Lin *et al*., 2024), and VvWRKY8, that suppresses resveratrol biosynthesis in response to resveratrol accumulation (Jiang *et al*., 2019). Further, *GmJAZ1* genes are upregulated by ABA, whereas *GmWRKY72* is upregulated by WGE, suggesting that they may participate in distinct response pathways. The results may also inform on details of the canonical JA signaling pathway in soybean. Since the level of glyceollin biosynthesis elicited by WGE is increased by silencing *GmJAZ1* gene transcripts (Figure 4C), it is tempting to speculate that the degradation of GmJAZ1 proteins by the canonical JAZ signaling pathway is incomplete. In other words, reducing *GmJAZ1* transcripts would not increase JA signaling unless JAZ protein levels were further reduced by the gene silencing. The *P. sojae* Avh94 effector protein directly binds GmJAZ1/2 proteins to prevent their degradation, which suppresses JA signaling and enhances soybean’s susceptibility (Zhao *et al*., 2022), suggesting that the canonical JA signaling pathway is active in soybean and involved in providing incompatibility to *P. sojae*. Thus, the most parsimonious explanation may be that our results provide further support that the balance between JAZ protein degradation and *JAZ* gene transcription is important for determining the activation or repression JA signaling, at least in the context of GmJAZ1 and its regulation of phytoalexin biosynthesis.

Our RNA-seq analysis showed that both ABA signaling and *GmJAZ1* genes have opposite expression patterns compared to glyceollin biosynthetic genes (Figure 2B, 2C, Table 1). This raised the possibility that the ABA signaling and *GmJAZ1* genes function together in the same signaling pathway to suppress glyceollin biosynthesis. In support of this, ABA or dehydration treatments upregulated *GmJAZ1-8/9*, *GmJAZ1-1*, and *GmJAZ1-2* (Figure 3D). Importantly, *GmJAZ1-8/9*, *GmJAZ1-1*, and *GmJAZ1-2*, are the same genes that were silenced in all RNAi experiments that exhibited a derepression of glyceollin biosynthesis (Figures 3F, 3G, 4A, 4C). Specifically, *GmJAZ1-8/9* and *GmJAZ1-1* genes showed opposite expression patterns compared to glyceollin biosynthesis upon treatment with an ABA biosynthesis inhibitor (Figures 3B, 3C, 3D), suggesting that it is particularly these genes that endogenous ABA signaling is using to suppress glyceollin biosynthesis. It is perhaps noteworthy that the expression of those *GmJAZ1* genes is affected specifically by long-term (6 d) dehydration, ABA, and norflurazon treatments, not by short-term (24 h) ABA treatment (Figure 3D, Supplementary Figure S4). Since studies on the effects of ABA are typically conducted following shorter durations of ABA treatment (*i.e.* usually < 48 h), it is tempting to speculate that our long-term treatment may be why previous studies have, to our knowledge, not identified ABA as a positive regulator of *JAZ1* gene expression. Overall, the most parsimonious explanation of our results is a model by which dehydration stress activates ABA biosynthesis and signaling, which in turn upregulates the expression of select *GmJAZ1* genes that suppress glyceollin biosynthesis (Figure 7).

**Figure 7.**
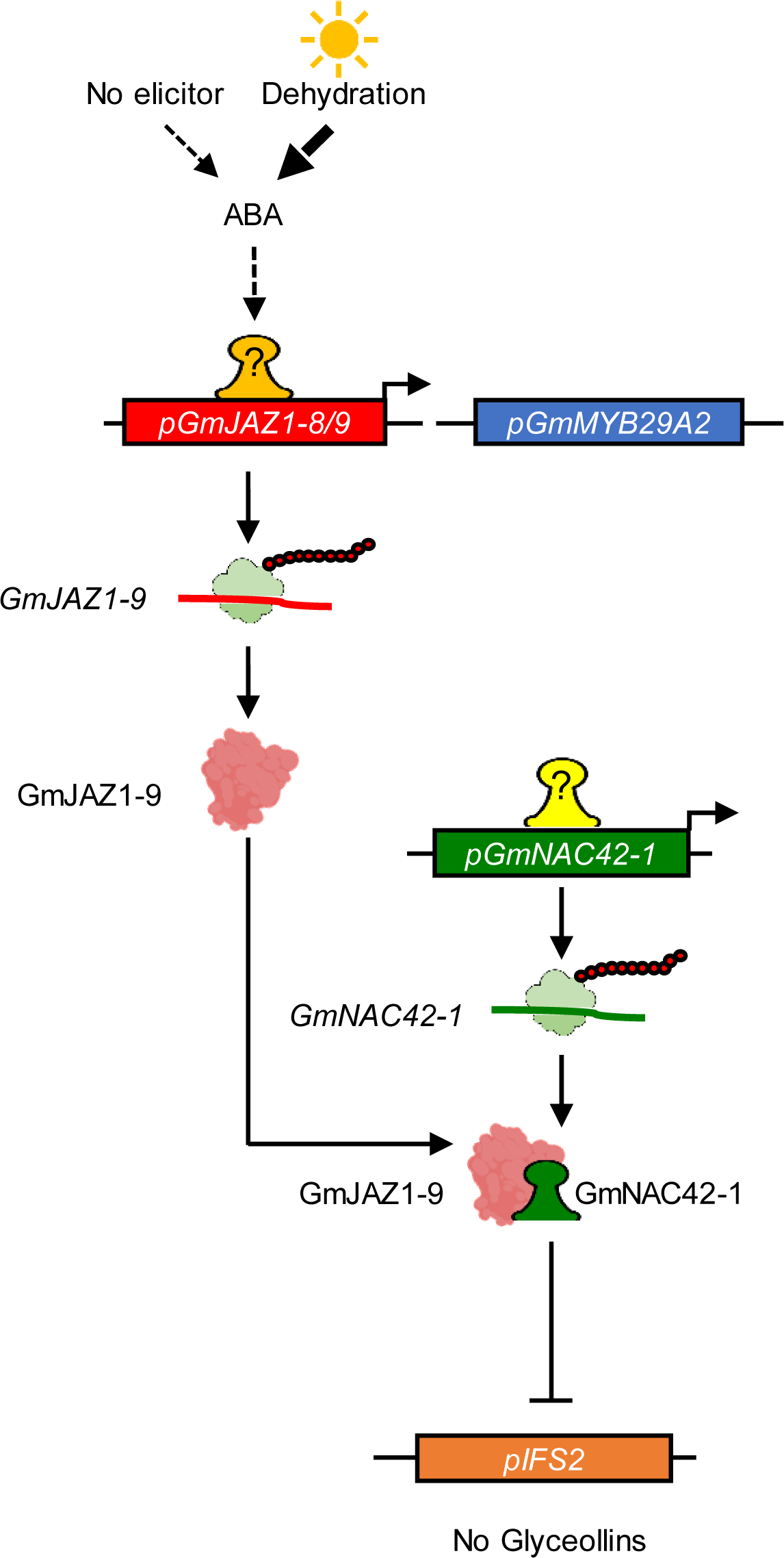
Schematic diagram of the negative regulation of glyceollin biosynthesis by the GmJAZ1-9-GmNAC42 protein-protein interaction. Drought induces ABA accumulation which stimulates *GmJAZ1* gene transcription *via* an unknown regulator. The resulting mRNA is translated and the GmJAZ1-9 protein physically interacts with GmNAC42 proteins to prevent them from activating glyceollin gene transcription. Broken arrows, multiple steps; solid arrows, single steps; colored boxes, gene promoters; bent arrows, gene transcription; brown globular structure, GmJAZ1-9 protein; colored bell-shaped structures, transcription factors.

Decades ago, ABA has been established as a negative regulator of phytoalexin biosynthesis, yet its molecular mechanism has remained unknown. Here, our experiments demonstrated that ABA treatment, or dehydration stress, which uses ABA as a prominent signal, upregulates the expression of select *GmJAZ1* genes. Based on the conventional JA signaling model, JAZ proteins negatively regulate JA signaling by physically interacting with transcription factor proteins to inhibit their activities in regulating gene transcription (Cheng *et al*., 2011, Fernández-Calvo *et al*., 2011, Qi *et al*., 2011, Sasaki-Sekimoto *et al*., 2014). Perhaps the most characterized JAZ target is MYC2, which is considered the ‘master’ regulator of JA defense responses (Kazan and Manners, 2013). Based on gene expression analyses and functional tests, MYC2 was found to be a positive regulator of insect defense, wound responses, flavonoid metabolism, and oxidative stress tolerance during JA signaling. However, MYC2 negatively regulates the expression of camalexin biosynthesis and microbial pathogen defenses. Since JA is a positive regulator of camalexin biosynthesis (Zhou *et al*., 2022), MYC2 is not a likely target of JA signaling for the activation of phytoalexin biosynthesis. Here, our experiments demonstrate that JAZ1 proteins from soybean, Arabidopsis, and grapevine, physically interact with their respective NAC42-type transcription factors by Y2H and BiFC assays (Figure 5B-5F). Notably, these NAC42-type proteins are transcriptional activators of their respective phytoalexin biosynthetic pathways (Saga *et al*., 2012, Vannozzi *et al*., 2018, Jahan *et al*., 2019). Our Y3H and promoter-reporter experiments demonstrate that the GmJAZ1-9-GmNAC42-1 interaction inhibits GmNAC42-1’s DNA binding activity and transactivation of glyceollin biosynthetic gene promoters (Figure 6A-6C). Taken together, our results support a working model whereby ABA-upregulated GmJAZ1 genes encode proteins that physically interact with NAC42-type transcription factors to inhibit their activation of phytoalexin biosynthesis (Figure 7). This model offers a plausible explanation as to why overexpressing *GmNAC42-1*, and/or its transcriptional activator GmMYB29A2, is insufficient to activate the expression of glyceollin biosynthetic genes in the absence of elicitation (Jahan *et al*., 2019, Jahan *et al*., 2020). Our future experiments will test whether simultaneously silencing or knocking-out *JAZ1* genes can be combined with overexpressing transcriptional activators, such as NAC42-type transcription factors, to constitutively activate phytoalexin biosynthesis. Since the Avh94 effector protein from *P. sojae* directly binds GmJAZ1/2 proteins, preventing their degradation, which suppresses JA signaling and enhances soybean’s susceptibility to *P. sojae* (Zhou *et al*., 2022), it is tempting to speculate that silencing select *GmJAZ1* genes in soybean could be exploited to enhance glyceollin-mediated resistance to microbial pathogens. In Arabidopsis, one-quarter of transgenic lines overexpressing JAZ1 lacking the Jas domain, that is required JAZ protein degradation, showed increased resistance to DC3000 infection (Thines *et al*., 2007).

## EXPERIMENTAL PROCEDURES

### Chemicals

Stocks of the ampicillin, kanamycin, gentamicin, hygromycin-B (50 mg/mL), and spectinomycin (50 mg/mL) were dissolved in distilled H2O. Stock MES hydrate (1 M) was adjusted to pH 5.6 using KOH. The stocks of acetosyringone (Cayman Chemical, MI, USA) and norflurazon were 100 mM in dimethyl sulfoxide and 5 mM in 10% ethanol (v/v), respectively. The stock of (+)-ABA and (+)-ABA-GE (Cayman Chemical and NRC Canada, respectively) were 40 mM in methanol. Glyceollin I and other isoflavone standards were purchased from Paul Erhardt (University of Toledo and Extrasynthese (France), respectively. Wall glucan elicitor (WGE) was isolated from *Phytophthora sojae* as described previously (Farrell *et al*., 2017). For an extensive list of manufacturers and catalog numbers of materials used in this study, see Supplementary Table S15.

### Maintenance of *P. sojae* and Isolation of WGE

Isolated *P. sojae* race 1 was a generous donation from Brett Tyler (Oregon State University). Pathogen maintenance was accomplished using a protocol previously described by (Jahan *et al*., 2020). Briefly, Williams 82 soybean seedlings were routinely inoculated with a mycelial plug (6–7 mm in diameter) and *P. sojae* was re-isolated to maintain pathogenicity. *P. sojae* was grown on lima bean culture plates (12-14 d) and then in liquid cultures for extraction of WGE according to (Farrell *et al*., 2017).

### Plant Materials and Growth Conditions

Williams 82 (W82) soybean plants were grown in the York University greenhouse and the seeds were collected in September. Soybean seeds were sterilized by 70% and 10% bleach as described by (Lin *et al*., 2023b). Sterilized seeds were grown 3 d under dark and 4 d under dim light (65 mEm^2^/s) on unbleached paper towels (Staples, ON, Canada) saturated with germination and co-cultivation (GC) medium. Soybean seedlings were grown in vermiculite at 22 °C under a 16 h photoperiod [cool white T5 fluorescent lights (250 mEm^2^/s)], and the first trifoliate leaf stage (∼ 8 days old) was used for dehydration, ABA, and norflurazon treatments (Jahan *et al*., 2019). *Nicotiana benthamiana* seedlings were used for the bimolecular fluorescence complementation (BiFC) assays. They were grown under the photoperiod of 16 h light 23 °C/8 h dark 21 °C (250 mEm^2^/s). Cucumber (*Cucumis sativus* variety Straight Eight, OSC Seeds, ON, Canada) were grown in vermiculite at 25 °C under a 12 h photoperiod [cool white T5 fluorescent lights (500 mEm^2^/s)].

### Cloning

The full length CDSs of GmJAZ1-9, GmMYB29A2, and GmNAC42-1 were PCR amplified from the cDNA of W82 hairy root treated with WGE (24 h) using specific primers (Table S16) and 2X Phusion Master Mix (ThermoFisher Scientific, MA, USA). The PCR amplicons were cloned into the entry vector pDONR221 using BP clonase (Invitrogen, ON, Canada). For overexpressing GmJAZ1-9 into hairy roots, entry clone (GmJAZ1-9-pDONR221) were LR (Invitrogen, ON, Canada) recombined downstream of the CaMV 35S promoter in the destination vector pGWB2. For subcellular localization assessment, entry clone were LR recombined downstream of GFP in the pGWB6 vector. For RNAi silencing, a 302-bp region corresponding to the CDS of GmJAZ1-9 was PCR amplified from cDNA, BP recombined into pDONR221, and after sequencing was LR subcloned into the RNAi destination vector pANDA35HK. For yeast hybrid assays, entry vectors were LR recombined downstream of the GAL4 activation domain and the GAL4 binding domain in the pDEST-GADT7 and pBD-Gal4-GW-C1 and pDR-GW vectors for expression in yeast for Y1H, Y2H, and Y3H, respectively. The CDSs of *GmJAZ1-9, GmMYB29A2, GmNAC42-1, AtJAZ1, AtNAC042, VvJAZ9, VvNAC36* without stop codon were amplified by PCR and BP cloned into pDONR221, then LR recombined into the BiFC destination vectors pB7WGYN9 and pB7WGYC9 (kindly provided by Dr. Zhao, UTSC). For protoplast transfection, entry vectors harboring the CDSs were LR recombined downstream of the CaMV 35S promoter in the destination vector p35S:HA-GW and entry vectors of the *pPTS1* and *pHIDH* promoters were LR recombined upstream of the *Luciferase* gene in the destination vector pBT10.

### Hairy Root Transformation

Soybean cotyledons were transformed with *Agrobacterium rhizogenes* strain K599 containing the empty vectors, overexpression, or RNAi constructs using our recently published protocol (Lin *et al*., 2023b). Briefly, *A. rhizogenes* (strain K599) harboring the plasmid of interest resuspended in phosphate buffer (pH = 7.5) containing 100 μM acetosyringone, OD_600_ was adjusted to 0.5-0.8 using a spectrophotometer (VE-5000V, Velab, TX, USA). Soybean cotyledons were infected by the agrobacteria suspension through three cuts on the adaxial surface. After three days co-culture on the filter paper saturated with GC media under a 16 h photoperiod (∼65 mEm^2^/s) and then, transferred into hairy root growth (HRG) medium containing timentin (500 mg/L Caisson Laboratories, UT, USA). After 3 weeks, transgenic secondary hairy roots were harvest and treated with WGE and H2O as described by (Lin *et al*., 2024).

### Composite plant transformation

The production of soybean composite plants was performed as described by (Kereszt *et al*., 2007). Briefly, soybean seeds were sterilized using a solution of 30% (w/w) hydrogen peroxide/96% (v/v) ethanol/sterilized distilled H2O (1:7.5:10). The seeds were then evenly distributed onto sterilized vermiculite that had been saturated with sterilized water. The transparent container lids were sprayed with 75% ethanol and allowed to dry. Seeds were left to germinate for five days. On the fifth day post-germination, a 1 mm needle was used to scrape *A. rhizogenes* strain K599, which harbored either the empty vector GUS or the JAZ1 silencing construct, from LB agar plates. The bacterial mass was used to pierce the hypocotyls of the soybean seedlings by gently pushing the needle tip into the seedlings. The transformed seedlings were placed in a growth chamber set at 28 °C with a 12-hour photoperiod, provided by cool white T5 fluorescent lights (250 mEm^2^/s), and maintained at 25 °C in darkness during the night. The infected soybean seedlings and transparent lid of the growth chamber was sprayed with 6X diluted MS medium daily to maintain humidity. Approximately three weeks after transformation, transgenic hairy roots were transplanted into soil and allowed to acclimate for one week before undergoing dehydration treatment.

### Stress Treatments and Elicitation

Stress treatments (dehydration and acidity) and WGE elicitation were performed following the protocols previously reported by (Jahan *et al*., 2019, Jahan *et al*., 2020) with slight modifications. Briefly, five soybean seedlings were wrapped with cascades single-fold paper towel saturated with MS medium for all stress treatments. For acidity stress, the wrapped seedlings were placed into a 100 mL beaker containing 50 mL of MS medium and then inserted inside a sterile 1000 mL beaker (contained 100 ml sterilized H2O). Another 1000 mL beaker was put on top upside down sealed with parafilm. For acidity stress with ABA treatment, the 50 mL of MS medium [pH 3.0] contained 100 µM ABA. For dehydration stress, the medium-saturated paper towel was dry gradually for 6 d. For dehydration stress and norflurazon treatment, the wrapped seedlings were submerged 50 mL of MS medium containing 10 µM norflurazon, incubated for 2 h, and then dry in the 100 mL beaker. For hairy roots elicitation, the transgenic secondary roots were cut into 1-cm pieces and placed on HRG plates overlaid with 80 µL of WGE (20 mg/mL), ABA (100 µM), WGE-ABA (20 mg/mL, and 100 mM, respectively) or H2O on top of the HRs piles, respectively. The hairy roots were then incubated for 24 h under 500 mEm^2^/s. Treated plant materials were used directly for metabolite extractions or flash frozen and lyophilized (BenchTop Pro, SP Scientific, PA, USA) for RNA extraction.

### ABA and ABA-GE Analysis

Hormones were extracted using a method adapted from (Forcat *et al*., 2008). Briefly, freeze-dried seedlings powder (5 mg) was extracted with 400 µl of a solvent mixture (methanol:acetic acid:H2O = 10:1:89) using stainless steel beads in a Mixer Mill MM400 (Retsch, CT, USA). The sample was pre-frozen for 3 min, followed by 30 min on ice. To ensure accurate quantification, 5µM (+)-catechin was used as an internal standard (Kovinich *et al*., 2011). The mixture was centrifuged to separate the supernatant, which was then pooled and concentrated by lyophilization. The dried residue was reconstituted in 100 µL of the same solvent mixture and filtered through 0.2 µm PTFE filters prior to LC-MS analysis. Hormone analysis was performed using a Q-Exactive Orbitrap mass spectrometer (Thermo Fisher Scientific, MA, USA) coupled with an Accela UHPLC (Thermo Fisher Scientific, MA, USA) system. Separations were achieved on an Acquity UPLC BEH Shield RP18 column (130Å, 1.7 µm) with the gradient reported by (Farrell *et al*., 2017). For mass spectrometry, Selected Ion Monitoring (SIM) mode was used with specific parameters for each compound. The dwell time was 10 msec. The Q1 mass, Q3 mass, declustering potential (DP), collision energy (CE), and collision exit potential (CXP) were specific for each compound, the parameters used were ABA Q1 263 Q3 153 DP −95 CE −18 CXP −17, ABA-GE Q1 425.2 Q3 153 DP −140 CE −32 CXP −53, and (+)-catechin Q1 288.8 Q3 125.1 DP −50 CE −10 CXP −13. Data normalization was performed based on the internal standard and dry tissue weight using Xcalibur software and Excel.

### Isoflavonoid Analyses

Fresh tissue of elicited hairy roots (90-110 mg; 24 h) was transferred in a 2 mL micro-centrifuge tube and placed on ice, following the extraction protocol according to (Farrell *et al*., 2017). Briefly, Tissues were ground for 3 minutes using 5 mm stainless steel balls in an MM400 mixer mill (Retsch, CT, USA) at 30 Hz. After adding 80% ethanol (1 µL per mg of tissue), the mixture was centrifuged at 17,000×g for 3 minutes. The supernatant was clarified at −20 °C overnight, then centrifuged again. The clear supernatant was filtered through a 0.2 µm filter, and 1 µL of this filtrate was analyzed with a Vanquish UHPLC system (Thermo Scientific, MA, USA) featuring a quaternary pump, autosampler, and Diode Array Detector (DAD). Isoflavonoid levels were quantified using Chromeleon 7.2.10 software, following the procedure outlined by (Jahan *et al*., 2020).

### RNA Extraction and Gene Expression Measurements

Total RNA from soybean tissue was extracted using the HiPure Total RNA mini Kit (GeneBio System, ON, Canada) following the manufacturer’s protocol as described previously (Lin *et al*., 2024). cDNA was synthesized using a DNA synthesis kit (GeneBio System, ON, Canada) following the manufacturer’s instructions. Quantitative RT-PCR was performed on a Bio-Rad CFX96 machine (Bio-Rad Laboratories, ON, Canada) as described by (Lin *et al*., 2024). Briefly, reactions (10 μL) contains 2 μL diluted cDNA (10X), 0.8 μL of 5mM forward and reverse primers (Supplemental Table S16), 2.2 μL RNAse free H2O, 5 μL GB-Amp^TM^ Sybr Green qPCR mix (GeneBio System, ON, Canada). The thermal cycling conditions were: initial denaturation at 95 °C for 3 minutes, followed by 40 cycles of 95 °C for 5 seconds and 60 °C for 30 seconds, with a melt curve analysis from 65 °C to 95 °C. GmUbiquitin3 was used as the internal reference for transcript normalization. Data were analyzed using the comparative CT method: expression = 2^-[Ct(gene) - Ct(UBIQUITIN3)].

### RNA-Seq and other in Silico Analyses

RNA sequencing was performed as described by (Jahan *et al*., 2019). Total RNA was extracted from powdered, lyophilized soybean seedlings using the Spectrum Plant Total RNA Kit (Sigma-Aldrich, MO, USA). We prepared 12 RNA libraries from three seedlings per stress treatment and their controls. The libraries were prepared by the Genomics Core Facility at West Virginia University. RNA quality was assessed with an RNA Nano 6000 Chip on an Agilent 2100 Bioanalyzer, and only samples with an RNA Integrity Number (RIN) greater than 8.0 were used. RNA (750 ng) was converted to cDNA using a stranded library prep kit (KAPA Biosystems, MA, USA) with nine PCR cycles. The cDNA libraries were quantified using a Qubit and pooled in equal amounts for sequencing at Marshall University Genomics Core on an Illumina HiSeq1500 system, generating 100 bp paired-end reads with eight libraries per lane in high-output mode. To ensure analysis robustness, principal component analysis, heatmap visualization, and clustering were performed. Gene homologs were identified using reciprocal BLASTP comparisons. Gene ontology analysis was carried out using the SoyBase Gene Model Data Mining and Analysis tool (https://www.soybase.org/goslimgraphic_v2/dashboard.php). Protein clustering was done with MEGA 11.0 software using MUSCLE for sequence alignment.

### Subcellular Localization

Soybean hairy roots transformed with nlsGFP-pGWB6 (positive control), pGWB2 (negative control), or nlsGFP-GmJAZ1-9-pGWB6 were harvested and stained with 4’,6-diamidino-2-phenylindole (DAPI, 6 μg/ml, from a 6 mg/mL stock in DMSO) according to the manufacturer’s instructions (Cayman Chemical, MI, USA). Three-to-four roots per genotype per two independent transformation events were analyzed. Confocal images were acquired using an LMS 700 Confocal Laser Scanning Microscope (Zeiss, Jena, Germany) with an X-Cite 120 fluorescence lamp, filter set for DAPI, and Zen Black software. Imaging was performed using Zen Blue software. Excitation and emission spectra were 488 nm and 500–550 nm for GFP and 405 nm and 358–461 nm for DAPI, respectively.

### BiFC Assays

*Agrobacterium tumefaciens* strain GV3101 was transformed with BiFC constructs and cultured overnight in the infiltration media (10 mM MES-KOH, 100 μM Acetosyringone, 50 μg/ml Spectinomycin, 50 μg/ml Gentamycin). After resuspending the *A. tumefaciens* in solution (10 mM MES, 3.2 g/L LB (Bioshop, ON, Canada) + 1000X Vitamins mixture (Caisson Laboratories, UT, USA), 20 g/L sucrose (Bioshop, ON, Canada), and 200 μM Acetosyringone), 4-5 weeks *N. benthamiana* seedlings were subjected to infiltration with BiFC construct pairs simultaneously. The epidermis was peeled using nail polish and imaged after 36 h. *N. benthamiana* epidermal peels were analyzed using a confocal microscope (LMS 700, Carl Zeiss, Jena, Germany) under a bright field and YFP (excitation/emission wavelength=514 nm/525-552 nm).

### Protoplast Transactivation Assay

Protoplast transactivation assay was performed according to the protocol described by (Wehner *et al*., 2011). Protoplast from *Cucumis sativus* was obtained by digestion of 6-day old leaves with cellulase for 8 hours and by density gradient separation (40% sucrose - 10% mannitol). The isolated protoplasts were transformed with both *pBT10-promoter* and *p35S:HA-TRANSCRIPTION FACTOR* constructs simultaneously, followed by incubation for 18 hours at 25 °C in dark. Luminescence measurements was performed using Dual-Luciferase Reporter Assay System protocol (Promega, WI, USA) and the multi-detection microplate reader (Agilent, CA, USA)

### Yeast Hybrid Assays

For yeast one-hybrid, yeast (*Saccharomyces cerevisiae*) strain YM4271 was co-transformed with both TF-pDEST-GADT7 and Promoter-pMW#2. Transformants were selected on media without amino acids Leucine and Histidine (SD-Leu-His). Positive Protein-DNA interactions (PDIs) were selected by growth on SD-Leu-His plates that contained 20 mM 3-amino-1,2,4-triazole (3AT; Fisher Scientific, MA, USA) as described (Jahan *et al*., 2019). For all yeast hybrid assays, representative results of three biological replicates, with each biological replicate derived from independent transformation events (colonies).

For yeast two-hybrid, *S. cerevisiae* strain PJ69-4a were simultaneously transformed with both pDEST-GADT7-prey and pBD-Gal4-GW-C1-bait (Supplementary Table S17). Transformants were selected on media lacking Leu, and Trp (SD-Leu-Trp) and then confirmed by colony PCR. Autoactivation was tested for yeast PJ69-4a strains transformed with pBD-Gal4-GW-C1-bait and empty pDEST-GADT7 on SD-His-Leu-Trp, and positive protein-protein interactions were determined by growth in SD-His-Leu-Trp medium that contained 10 mM 3AT.

For yeast three hybrid, the GmJAZ1-9-pDR construct or the empty vector pDR-GW was transformed into yeast that was previously transformed with the soybean transcription factor constructs GmNAC42-1-Gal4AD or GmMYB29A2-Gal4AD and the promoters *pHIDH* and *pPTS1* using the lithium acetate method. Transformants were selected on media without Leu, His, and Uracil (SD-Leu-His-Ura). Positive PDIs were selected by growth on SD-Leu-His-Ura plates that contained 20 mM 3AT.

### Statistics

The Tukey post hoc test and one-way single factor ANOVA were applied to determine whether any statistically significant differences existed between group means between treatments and genotypes. As mentioned in the figure legends, different letters show significant differences by single factor ANOVA, Tukey post hoc test (*P* < 0.05, α = 0.05). ANOVA values were calculated using the Data Analysis function of Excel (Microsoft) and Tukey values were using the formula M1 − M2 / (√(MS_w_(1/n)), where M1 = treatment/group mean 1, M2 = treatment/group mean 2, MS_w_ = mean square error within group, and n = sample size of each group. Paired comparisons were calculated using students *t*-test in Excel software, asterisks indicate significant differences (*P* < 0.05).

### Accession Numbers

GmJAZ1-1, Glyma.11G038600; GmJAZ1-2, Glyma.01G204400; GmJAZ1-3, Glyma.09G071600; GmJAZ1-4, Glyma.16G081800; GmJAZ1-5, Glyma.08G264700; *GmJAZ1-6*, Glyma.15G179600; GmJAZ1-7, Glyma.04G013800; GmJAZ1-8, Glyma.17G047700; *GmJAZ1-9*, Glyma.13G112000; *GmMYB29A2*, Glyma.02G005600; *GmNAC42-1*, Glyma.02G284300; HIDH, Glyma.01G239600; PTS1, Glyma.19G151100; AtJAZ1, AT1G19180; AtNAC042, AT2G43000; VvJAZ9, VIT_211s0016g00710; VvNAC36, VIT_12s0028g00860.

## Acknowledgments

The authors gratefully acknowledge Baodong Wu for his expertise in cultivating the plants. Nora Wehner (Julius-von-Sachs-Institut) for the pBT10 and p35S:HA-GW vectors, the Arabidopsis Biological Resource Center (ABRC) for the *ANAC042* clone and the yeast hybrid vectors, Karl Andrew White (York University) for the cucumber seeds and protoplast transfection protocol, Erich Grotewold (Michigan State University) for the vectors pDEST-GADT7, pBD-Gal4-GW-C1, pDR-GW, and Rongmin Zhao (University of Toronto Scarborough) for the vectors pB7WGYN9 and pB7WGYC9.

## Short Legends for Supporting Information

### Supporting Figures

Figure S1. Repeat of experiment shown in Figure 2D.

Figure S2. Repeat of experiment in Figure 3B.

Figure S3. Repeat of experiment in Figure 3C.

Figure S4. Expression levels of ABA-responsive genes in W82 seedlings treated with 100 µM ABA for 24 h measured by qRT-PCR.

Figure S5. Repeat the hairy root experiments for Figures 4A, 4C, 4E, 4F.

Figure S6. Additional images and negative controls for the BiFC experiments shown in Figures 5C and 5F.

Supporting Tables:

Table S1. Statistical results for metabolite measurements shown in Figure 2D and Figure S1.

Table S2. Statistical results for metabolite measurements shown in Figure 3A.

Table S3. Statistical results for metabolite measurements shown in Figure 3B and Figure S2.

Table S4. Statistical results for metabolite measurements shown in Figure 3C and Supplementary Figure S3.

Table S5. Statistical results for gene expression measurements shown in Figure 3D.

Table S6. RNAi trigger counts in potential gene targets.

Table S7. Statistical results for gene expression measurements shown in Figures 3F.

Table S8. Statistical results for metabolite measurements shown in Figure 3G.

Table S9. T-test values of gene expressions changes for RNAi-GmJAZ1 versus RNAi-vector shown in Figure 4A.

Table S10. T-test values of gene expressions changes for RNAi-GmJAZ1 versus RNAi-vector shown in Figure 4B.

Table S11. T-test values of total glyceollins metabolite changes for RNAi-GmJAZ1 versus RNAi-vector shown in Figure 4C.

Table S12. T-test values of total glyceollins metabolite changes for RNAi-GmJAZ1 versus RNAi-vector shown in Figure 4D.

Table S13. T-test values of gene expressions changes for RNAi-GmJAZ1 versus RNAi-vector shown in Figure 4E.

Table S14. T-test values of total glyceollins metabolite changes for RNAi-GmJAZ1 versus RNAi-vector shown in Figure 4F.

Table S15. Manufacturers of materials used in this study and item catalog numbers.

Table S16. Oligonucleotides used in this study.

Table S17. Y2H constructs.

## Author Contributions

JL, IM (Ivan Monsalvo), ML, AJ, DW, and IM (Izabella Martirosyan) conducted the experiments. JL, AJ, IM (Ivan Monsalvo), ML and NK analyzed the data. NK conceived of the study and wrote the manuscript with input from all other authors. All authors read and approved of its content.

## Data Availability Statement

The RNA-seq datasets analyzed in this study were published in (Jahan *et al*., 2019) and can be found in the Gene Expression Omnibus (https://www.ncbi.nlm.nih.gov/geo/) under the series accession GSE112584.

