## Supplemental Figures for "ABA-regulated JAZ1 Proteins Bind NAC42 Transcription Factors to Suppress the Activation of Phytoalexin Biosynthesis in Plants": Figure S1. Metabolite amounts - Repeat of Figure 2D.pptx

### Slide 1
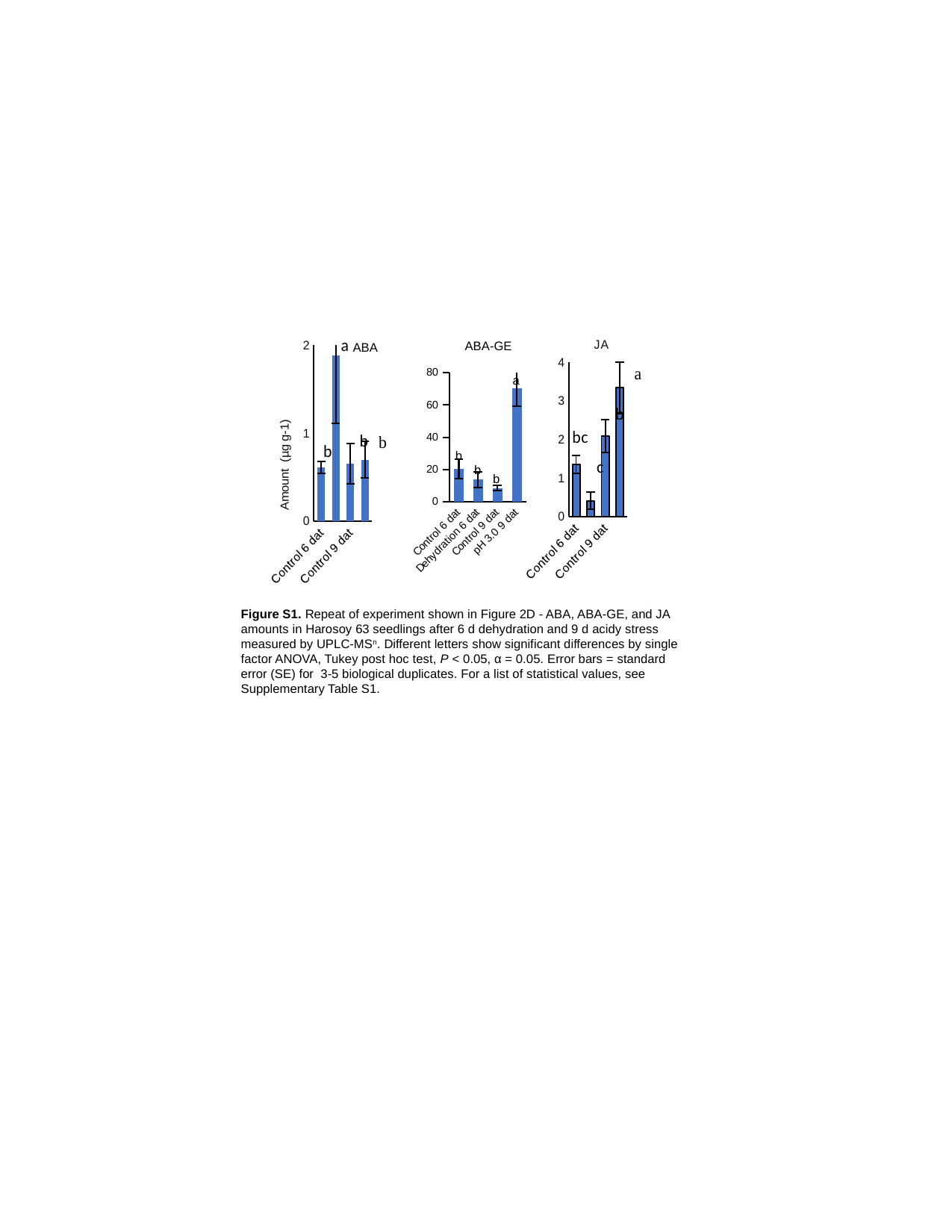

#### Chart
| Category | |
|---|---|
| Control 6 dat | 0.6087638584956556 |
| Dehydration 6 dat | 1.883082347032672 |
| Control 9 dat | 0.6531238140482172 |
| pH 3.0 9 dat | 0.6954065140282032 |
#### Chart: JA
| Category | |
|---|---|
| Control 6 dat | 1.3416659857982292 |
| Dehydration 6 dat | 0.4028470358633865 |
| Control 9 dat | 2.08066515052 |
| pH 3.0 9 dat | 3.3327048315999996 |ABA-GE
ABA
#### Chart
| Category | |
|---|---|
| Control 6 dat | 20.54823902174844 |
| Dehydration 6 dat | 13.820776287224476 |
| Control 9 dat | 8.605638190794563 |
| pH 3.0 9 dat | 70.40950701678862 |a
b
b
b
Figure S1. Repeat of experiment shown in Figure 2D - ABA, ABA-GE, and JA amounts in Harosoy 63 seedlings after 6 d dehydration and 9 d acidy stress measured by UPLC-MS^n^. Different letters show significant differences by single factor ANOVA, Tukey post hoc test, P < 0.05, α = 0.05. Error bars = standard error (SE) for 3-5 biological duplicates. For a list of statistical values, see Supplementary Table S1.
