## Supplemental Figures for "ABA-regulated JAZ1 Proteins Bind NAC42 Transcription Factors to Suppress the Activation of Phytoalexin Biosynthesis in Plants": Figure S2. Metabolite amounts - Repeat of Figure 3B.pptx

### Slide 1
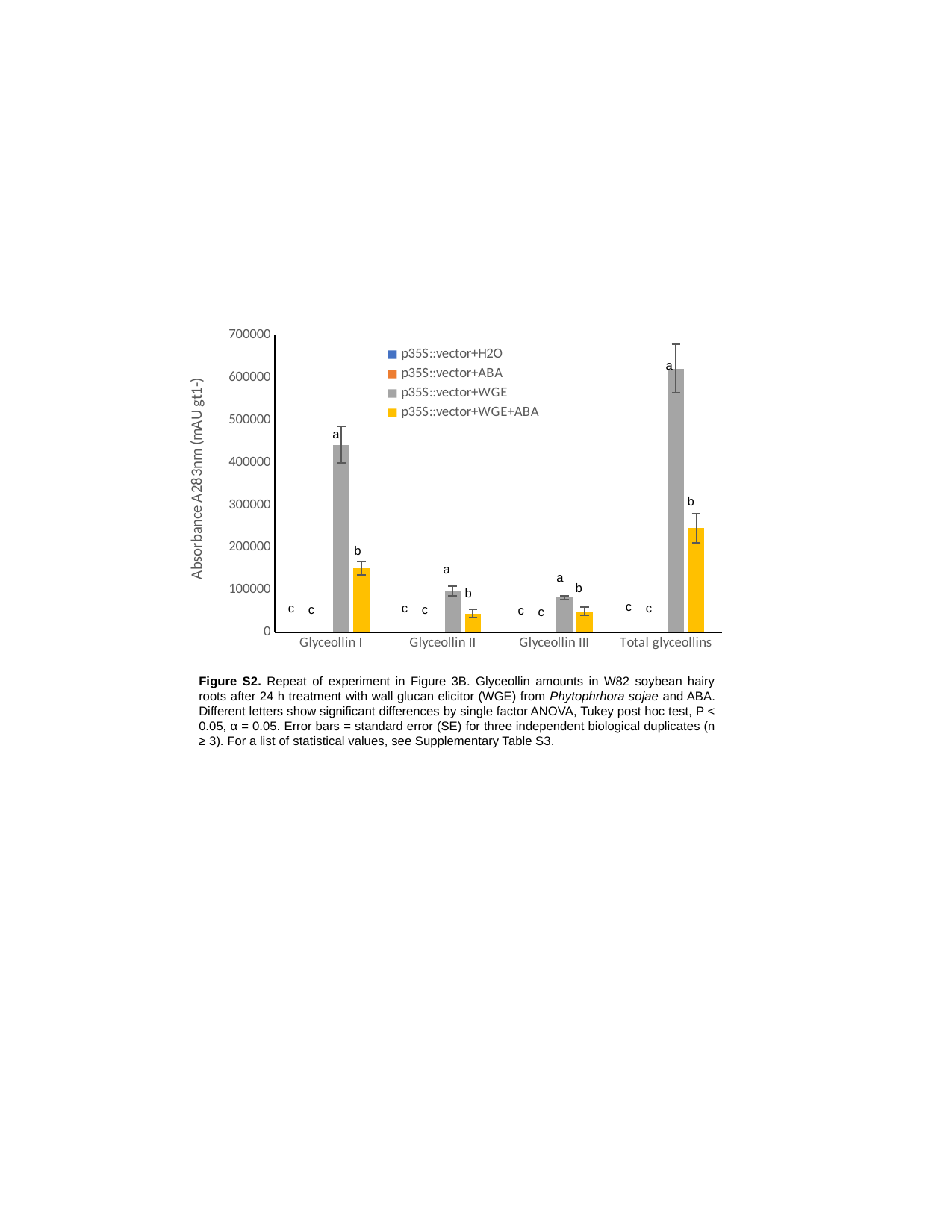

#### Chart
| Category | p35S::vector+H2O | p35S::vector+ABA | p35S::vector+WGE | p35S::vector+WGE+ABA |
|---|---|---|---|---|
| Glyceollin I | 0.0 | 0.0 | 442112.29500000004 | 151406.0 |
| Glyceollin II | 0.0 | 0.0 | 97664.09 | 44476.455 |
| Glyceollin III | 0.0 | 0.0 | 81794.56999999999 | 49452.62 |
| Total glyceollins | 0.0 | 0.0 | 621570.955 | 245335.07499999995 |a
a
b
b
a
a
b
b
c
c
c
c
c
c
c
c
Figure S2. Repeat of experiment in Figure 3B. Glyceollin amounts in W82 soybean hairy roots after 24 h treatment with wall glucan elicitor (WGE) from Phytophrhora sojae and ABA. Different letters show significant differences by single factor ANOVA, Tukey post hoc test, P < 0.05, α = 0.05. Error bars = standard error (SE) for three independent biological duplicates (n ≥ 3). For a list of statistical values, see Supplementary Table S3.
