## Supplemental Figures for "ABA-regulated JAZ1 Proteins Bind NAC42 Transcription Factors to Suppress the Activation of Phytoalexin Biosynthesis in Plants": Figure S3. Metabolite amounts - Repeat of Figure 3C.pptx

### Slide 1
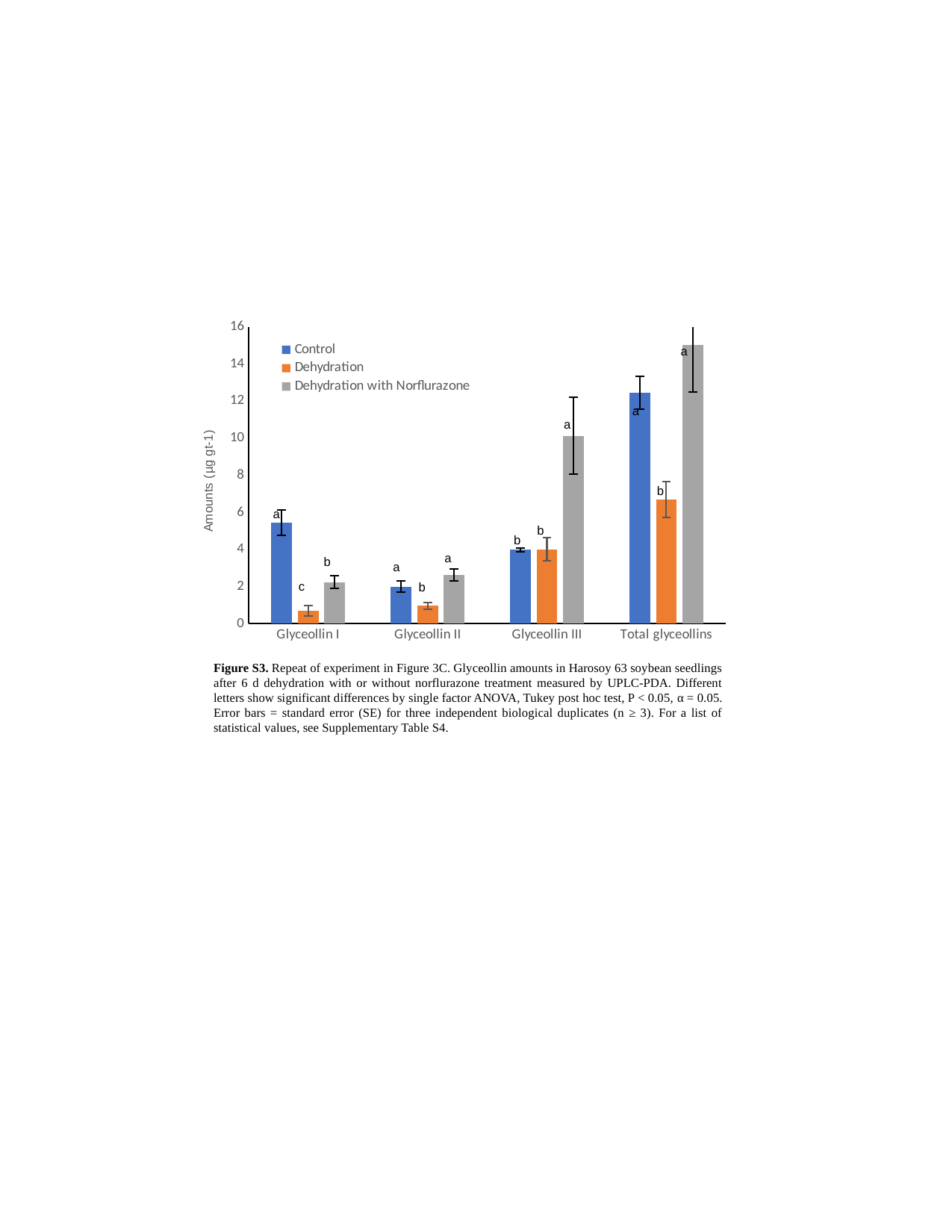

#### Chart
| Category | | Dehydration | Dehydration with Norflurazone |
|---|---|---|---|
| Glyceollin I | 5.4341382933333335 | 0.6897253333333335 | 2.246888 |
| Glyceollin II | 2.00625552 | 0.9795356800000001 | 2.6263720000000004 |
| Glyceollin III | 3.98824144 | 4.014208320000001 | 10.127262666666669 |
| Total glyceollins | 12.439635253333334 | 6.694469333333334 | 15.01152266666667 |a
a
a
b
a
b
b
a
b
a
c
b
Figure S3. Repeat of experiment in Figure 3C. Glyceollin amounts in Harosoy 63 soybean seedlings after 6 d dehydration with or without norflurazone treatment measured by UPLC-PDA. Different letters show significant differences by single factor ANOVA, Tukey post hoc test, P < 0.05, α = 0.05. Error bars = standard error (SE) for three independent biological duplicates (n ≥ 3). For a list of statistical values, see Supplementary Table S4.
