## Supplemental Figures for "ABA-regulated JAZ1 Proteins Bind NAC42 Transcription Factors to Suppress the Activation of Phytoalexin Biosynthesis in Plants": Figure S4. Expression of ABA genes and GmJAZ1s at 24 h ABA treatment (earlier time point than Fig3D).pptx

### Slide 1
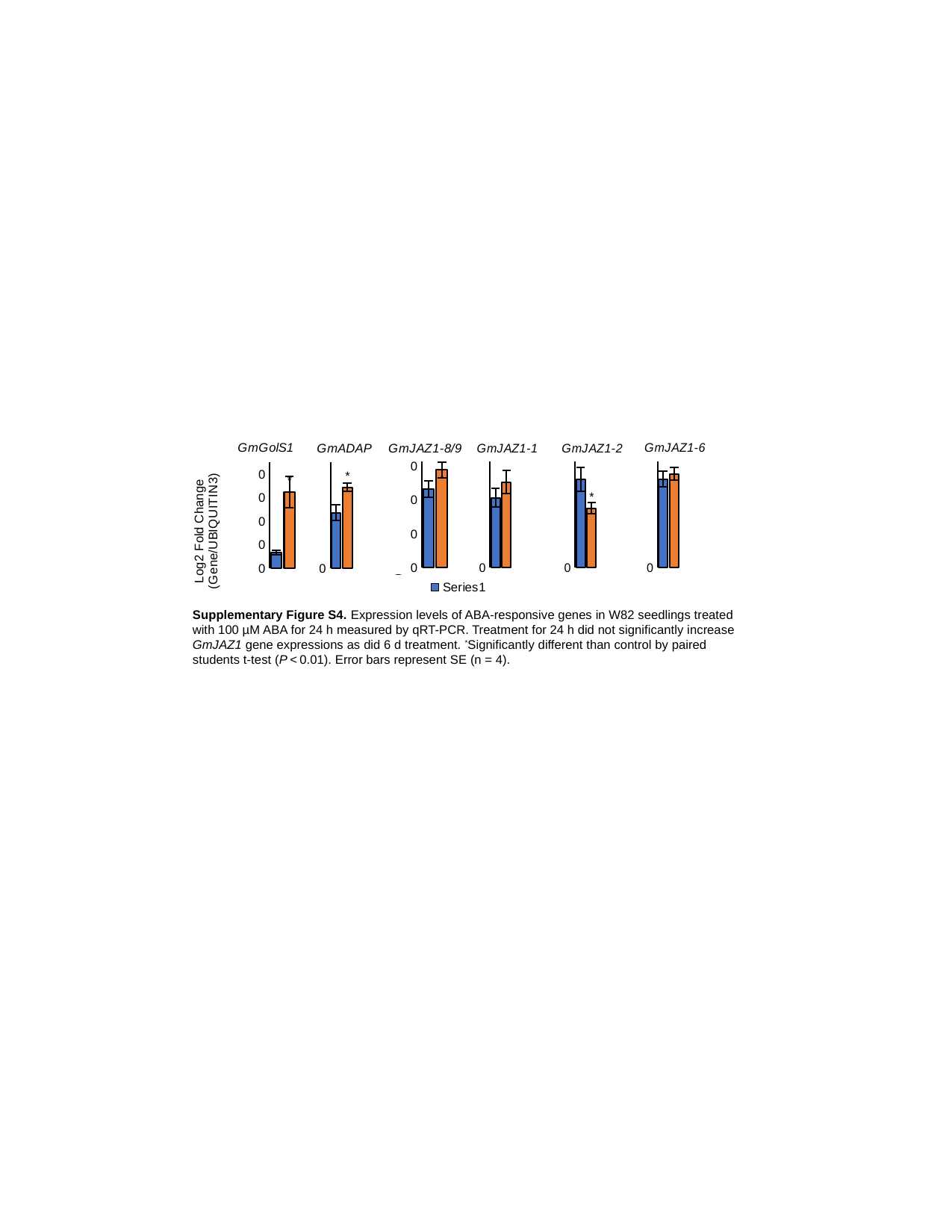

#### Chart: GmGolS1
| Category | |
|---|---|
| Control | 0.00026880324491392824 |
| ABA | 0.0012973184691493758 |
#### Chart: GmGolS1
| Category | |
|---|---|
| Control | 0.00026880324491392824 |
| ABA | 0.0012973184691493758 |
#### Chart: GmJAZ1-6
| Category | |
|---|---|
| H2O | 0.009592446708323965 |
| ABA | 0.01015603764143274 |
#### Chart: GmJAZ1-8/9
| Category | |
|---|---|
| H2O | 0.009299328130222127 |
| ABA | 0.011533711243367339 |
#### Chart: GmJAZ1-1
| Category | |
|---|---|
| H2O | 0.00043533797022137925 |
| ABA | 0.0005329550859077796 |
#### Chart: GmJAZ1-2
| Category | |
|---|---|
| H2O | 0.000208288292085526 |
| ABA | 0.0001400208025685376 |
#### Chart: GmADAP
| Category | |
|---|---|
| H2O | 0.00013095205872130542 |
| ABA | 0.00019069761037613842 |Supplementary Figure S4. Expression levels of ABA-responsive genes in W82 seedlings treated with 100 µM ABA for 24 h measured by qRT-PCR. Treatment for 24 h did not significantly increase GmJAZ1 gene expressions as did 6 d treatment. ^*^Significantly different than control by paired students t-test (P < 0.01). Error bars represent SE (n = 4).
