## Supplemental Figures for "ABA-regulated JAZ1 Proteins Bind NAC42 Transcription Factors to Suppress the Activation of Phytoalexin Biosynthesis in Plants": Figure S5. Repeat hairy roots experiments showing in Figure 4A 4C 4E 4F updated on Sep19 24.pptx

### Slide 1
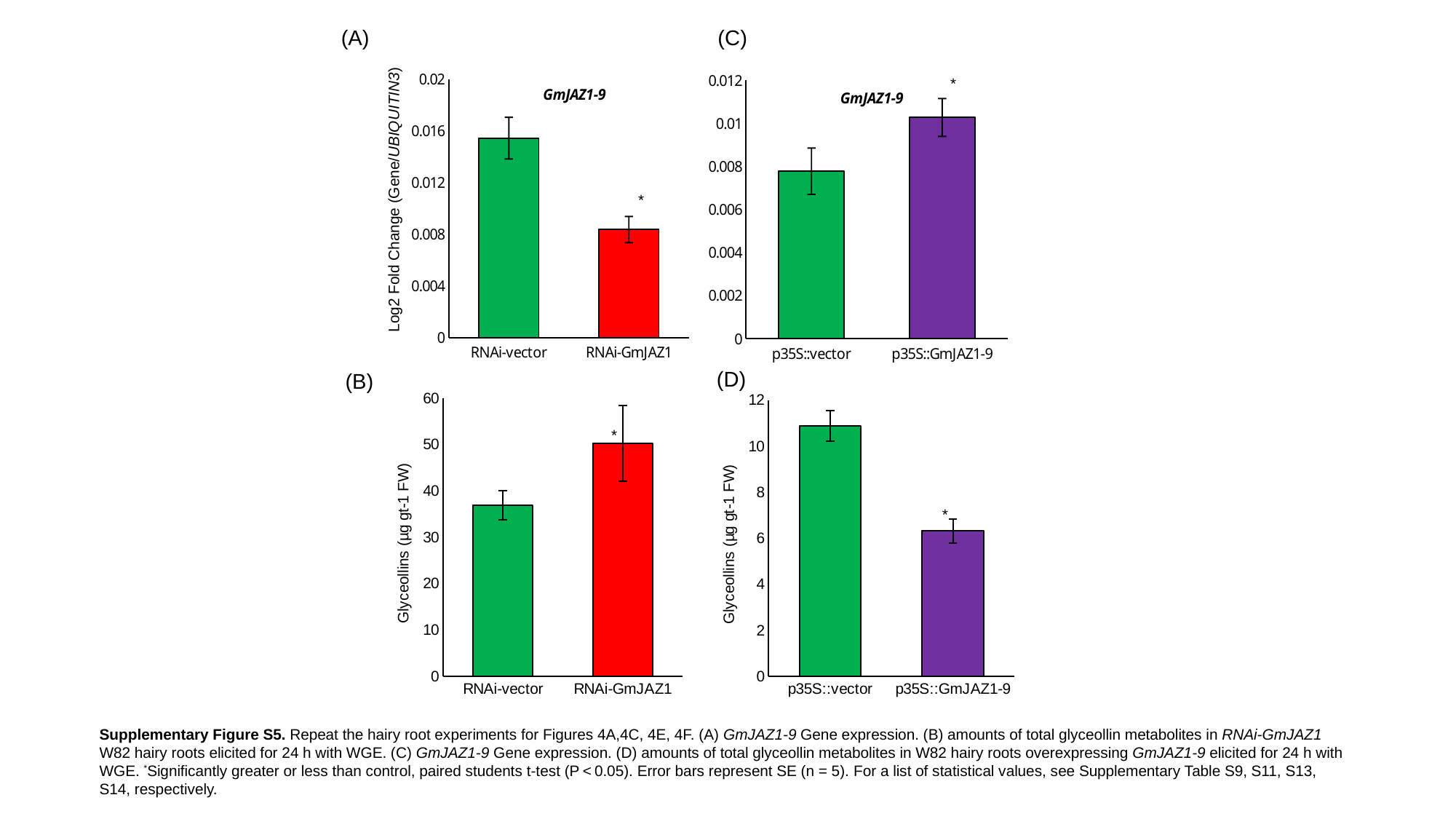

(A)
(C)
*
#### Chart: GmJAZ1-9
| Category | |
|---|---|
| RNAi-vector | 0.015429759099215143 |
| RNAi-GmJAZ1 | 0.008354161801420353 |
#### Chart: GmJAZ1-9
| Category | |
|---|---|
| p35S::vector | 0.007781787623258503 |
| p35S::GmJAZ1-9 | 0.010273679407404344 |*
Log2 Fold Change (Gene/UBIQUITIN3)
(D)
(B)
#### Chart
| Category | |
|---|---|
| RNAi-vector | 36.916242000000004 |
| RNAi-GmJAZ1 | 50.275654 |
#### Chart
| Category | |
|---|---|
| p35S::vector | 10.873434 |
| p35S::GmJAZ1-9 | 6.322976000000001 |*
*
Supplementary Figure S5. Repeat the hairy root experiments for Figures 4A,4C, 4E, 4F. (A) GmJAZ1-9 Gene expression. (B) amounts of total glyceollin metabolites in RNAi-GmJAZ1 W82 hairy roots elicited for 24 h with WGE. (C) GmJAZ1-9 Gene expression. (D) amounts of total glyceollin metabolites in W82 hairy roots overexpressing GmJAZ1-9 elicited for 24 h with WGE. ^*^Significantly greater or less than control, paired students t-test (P < 0.05). Error bars represent SE (n = 5). For a list of statistical values, see Supplementary Table S9, S11, S13, S14, respectively.
