## Supplemental Figures for "ABA-regulated JAZ1 Proteins Bind NAC42 Transcription Factors to Suppress the Activation of Phytoalexin Biosynthesis in Plants": Figure S6. Additional images and negative controls for BiFC assays Figure 5C 5F updated on Sep 19 24.pptx

### Slide 1
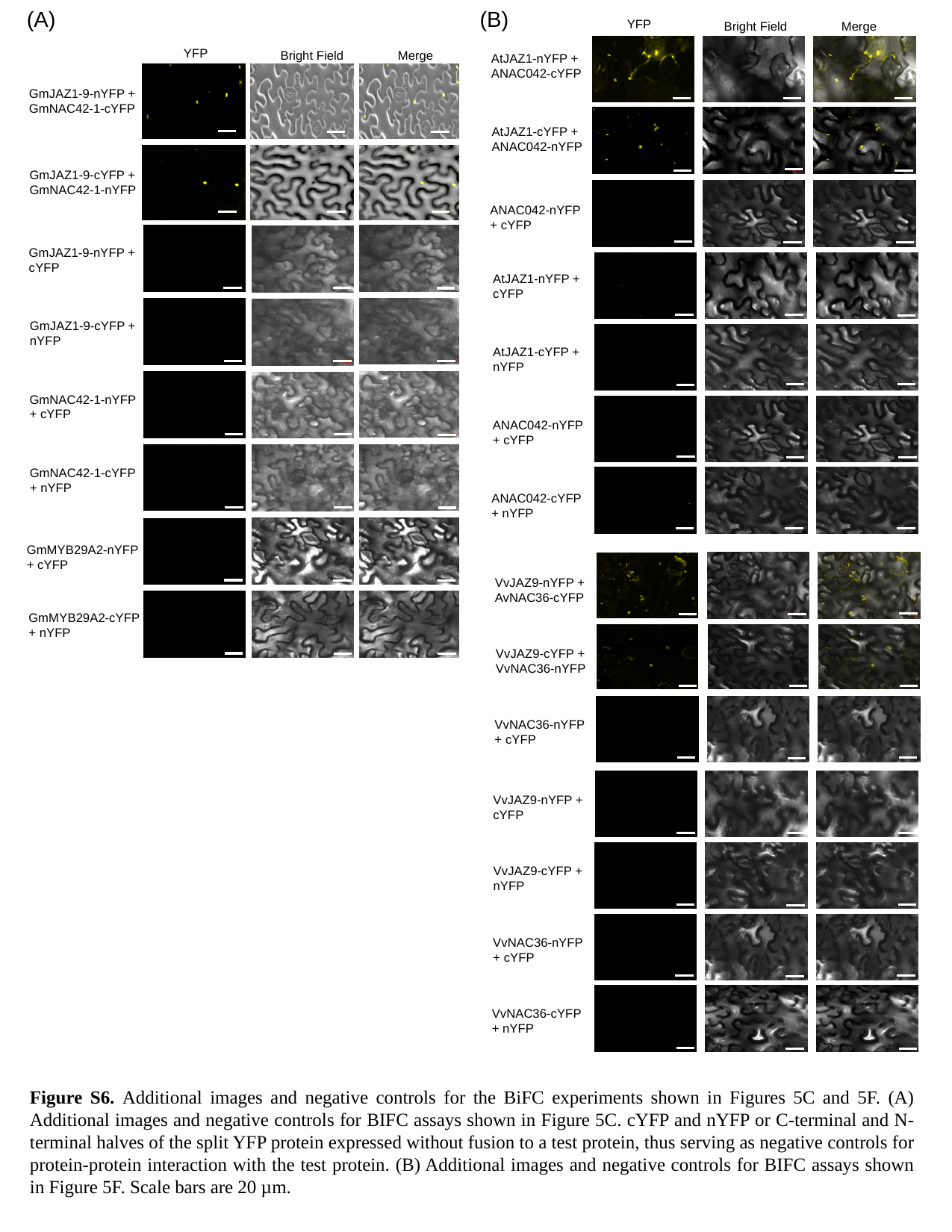

(A)
(B)
YFP
Bright Field
Merge
YFP
Bright Field
Merge
GmJAZ1-9-nYFP + GmNAC42-1-cYFP
GmJAZ1-9-cYFP + GmNAC42-1-nYFP
AtJAZ1-nYFP + ANAC042-cYFP
AtJAZ1-cYFP + ANAC042-nYFP
ANAC042-nYFP + cYFP
GmJAZ1-9-nYFP + cYFP
AtJAZ1-nYFP + cYFP
GmJAZ1-9-cYFP + nYFP
AtJAZ1-cYFP + nYFP
GmNAC42-1-nYFP
+ cYFP
ANAC042-nYFP + cYFP
GmNAC42-1-cYFP + nYFP
ANAC042-cYFP + nYFP
GmMYB29A2-nYFP + cYFP
VvJAZ9-nYFP + AvNAC36-cYFP
GmMYB29A2-cYFP + nYFP
VvJAZ9-cYFP + VvNAC36-nYFP
VvNAC36-nYFP + cYFP
VvJAZ9-nYFP + cYFP
VvJAZ9-cYFP + nYFP
VvNAC36-nYFP + cYFP
VvNAC36-cYFP + nYFP
Figure S6. Additional images and negative controls for the BiFC experiments shown in Figures 5C and 5F. (A) Additional images and negative controls for BIFC assays shown in Figure 5C. cYFP and nYFP or C-terminal and N-terminal halves of the split YFP protein expressed without fusion to a test protein, thus serving as negative controls for protein-protein interaction with the test protein. (B) Additional images and negative controls for BIFC assays shown in Figure 5F. Scale bars are 20 µm.
