## Supplemental Tables for "ABA-regulated JAZ1 Proteins Bind NAC42 Transcription Factors to Suppress the Activation of Phytoalexin Biosynthesis in Plants": Supplementary Table S16 - Oligonucleotides.pdf

**Table S16.** Oligonucleotides used in this study

| Oligo name | Sequence (5' to 3') | Usage |
|---|---|---|
| qUBf | GTGTAATGTTGGATGTGTTCCC | qRT-PCR |
| qUBr | CCGAACTTATTTTCAAGGGACAT | qRT-PCR |
| aNAC1f | AAAAAGCAGGCTTAACAATGGATGTGGCC AAGTTACA | Cloning |
| aNAC1r | AGAAAGCTGGGTGGCATGCACCCAAGAAA AGAG | Cloning |
| qNAC1f | CTCAGCAGCAGCACCAATAA | Colony PCR, qRT-PCR |
| qNAC1r | AAGTATGGCTTGCCGTTGTC | Colony PCR, qRT-PCR |
| MYB29f | GGGGACAAGTTTGTACAAAAAAGCAGGCT TAACAATGGTTAGAGCTCCTTGTTGTGA | Cloning |
| MYB29r | GGGGACCACTTTGTACAAGAAAGCTGGGT GTCAGAACTCTGGCAATTCGAT | Cloning |
| qMA2-3f | CTTGGACCCCAGAGGAAGA | Colony PCR, qRT-PCR |
| qMA2-3r | CAGCTCTTGCCACACCTTAAT | Colony PCR, qRT-PCR |
| JAZ1-6f | AAAAAGCAGGCTTAACAATGTCGAGCTCA TCGGAGTACT | Cloning |
| JAZ1-6r | AGAAAGCTGGGTGGCATTTTCACAACAAA AATAATGC | Cloning |
| cJAZ1-6f | CTCGGAATGACATCATGTGG | Colony PCR |
| cJAZ1-6r | GGTGCTCACGAATGGAGTTT | Colony PCR |
| qJAZ1-6f | TCATGTCCTATGCCACCAAA | qRT-PCR |
| qJAZ1-6r | AATGAAGGCTGGCTCTGTGT | qRT-PCR |
| JAZ1-9f | AAAAAGCAGGCTTAACAATGTCCAGCTCA TCGGAATATTT | Cloning |

| | | |
|---|---|---|
| JAZ1-9r | AGAAAGCTGGGTGTTAGACTTGTGTTGATTGAGCACC | Cloning |
| cJAZ1-9f | CCCCAGACTGCTTATCCTCA | Colony PCR |
| cJAZ1-9r | ATTCAGCCGGCTTACTTGAA | Colony PCR |
| siJAZ1-9f | AAAAAGCAGGCTGCTGCTCAGTTGACGATCTTT | RNAi cloning |
| siJAZ1-9r | AGAAAGCTGGGTGTTTATTTGATAGGGTGCTTTGG | RNAi cloning |
| qJAZ1-9f | GGTGCAGGCTTCAATTATTATG | qRT-PCR |
| qJAZ1-9r | GGGGAAAAGCAACAAATGAA | qRT-PCR |
| qJAZ1-1f | GGCAAGGGAATGTCTCAAAA | qRT-PCR |
| qJAZ1-1r | TGGCACTGGTTGGAATGATA | qRT-PCR |
| qJAZ1-2f | TTCCAGTTGTGTGAGCAAGG | qRT-PCR |
| qJAZ1-2r | ATGTCCTTGGCTTTGTCAGC | qRT-PCR |
| qJAZ4-1f | CCATCCGTACCTGCTCCTAA | qRT-PCR |
| qJAZ4-1r | TCCCAGCCAACAACATGATA | qRT-PCR |
| qHIDHf | GGAATCGATAACCCCTGGAT | qRT-PCR |
| qHIDHr | TGAACTCGTCTTTGCCAGTG | qRT-PCR |
| qPTS1f | GGCAAGGCACAAGGAATTTA | qRT-PCR |
| qPTS1r | ATGGTGCTTCCGTTGAACTC | qRT-PCR |
| qG4DTf | TGGCTTTCGGAGTTGGTATC | qRT-PCR |
| qG4DTr | GAACAGCATTTCCCATACCC | qRT-PCR |
| qIFS2f | GGAGAGGTTGTTGAGGGTGA | qRT-PCR |
| qIFS2r | GACCCTTGATGTGGTCCTTG | qRT-PCR |
| qC4Hf | AGGAGGAGTCGCTTGTTGAA | qRT-PCR |
| qC4Hr | CCCAAAGTGATGCCAAGAAT | qRT-PCR |
| RNAi-GmJAZ1-9 | GCTGCTCAGTTGACGATCTTTTATGCTGGGCAAGTTGTTGTGTTTGATGATTTTCCTGCTGAAAAATTGGAGGAGATAACGTCATTAGCCGGCAAGGGAATATCCCAAAGCCAAAACACCTCTGCTTATGCTCACACTCATAACCAGCAAGTGAATCATCCTTCCTTTGTTCCTAATATCTCCCCTCAAGCACCGTCCAGACCACTTGTTTGTGATCTGCCAATTGCTAGGAAAGCTTCACTTCATCGGTTCCTTTCTAAGAGAAAAGATAGAATTGCTGCCAAAGCACCCTATCAAATAAA | RNAi silencing trigger |
