## Supplemental Tables for "ABA-regulated JAZ1 Proteins Bind NAC42 Transcription Factors to Suppress the Activation of Phytoalexin Biosynthesis in Plants": Supplementary Table S17 - Y2H constructs.pdf

| Prey constructs | Bait Constructs |
|---|---|
| pDEST-GADT7-GmJAZ1-9 | pBD-Gal4-GW-C1-GmJAZ1-9 |
| pDEST-GADT7-GmNAC42-1 | pBD-Gal4-GW-C1-GmNAC42-1 |
| pDEST-GADT7-GmMYB29A2 | pBD-Gal4-GW-C1-GmMYB29A2 |
| pDEST-GADT7-AtJAZ1 | pBD-Gal4-GW-C1-AtJAZ1 |
| pDEST-GADT7-AtNAC042 | pBD-Gal4-GW-C1-AtNAC042 |
| pDEST-GADT7-AtMYB14 | pBD-Gal4-GW-C1-AtMYB14 |
| pDEST-GADT7- VvJAZ9 | pBD-Gal4-GW-C1-VvJAZ9 |
| pDEST-GADT7- VvNAC36 | pBD-Gal4-GW-C1-VvNAC36 |
| pDEST-GADT7-VVMYB14 | pBD-Gal4-GW-C1- VvMYB14 |
| pDEST-GADT7 | pBD-Gal4-GW-C1 |
